# Kar4, the Yeast Homolog of METTL14, is Required for mRNA m^6^A Methylation and Meiosis

**DOI:** 10.1101/2023.01.29.526094

**Authors:** Zachory M. Park, Abigail Sporer, Katherine Kraft, Krystal Lum, Edith Blackman, Ethan Belnap, Christopher Yellman, Mark D. Rose

## Abstract

*KAR4*, the yeast homolog of the mammalian mRNA N^6^A-methyltransferase complex component *METTL14*, is required for two disparate developmental programs in *Saccharomyces cerevisiae*: mating and meiosis. To understand *KAR4*’s role in yeast mating and meiosis, we used a genetic screen to isolate 25 function-specific mutant alleles, which map to non-overlapping surfaces on a predicted structure of the Kar4 protein (Kar4p). Most of the mating-specific alleles (Mat^-^) abolish Kar4p’s interaction with the transcription factor Ste12p, indicating that Kar4p’s mating function is through Ste12p. In yeast, the mRNA methyltransferase complex was previously defined as comprising Ime4p (Kar4p’s paralog and the homolog of mammalian METTL3), Mum2p (homolog of mammalian WTAP), and Slz1p (MIS), but not Kar4p. During meiosis, Kar4p interacts with Ime4p, Mum2p, and Slz1p. Moreover, cells lacking Kar4p have highly reduced levels of mRNA methylation during meiosis indicating that Kar4p is a key member of the methyltransferase complex, as it is in humans. Analysis of *kar4*Δ/Δ and 7 meiosis-specific alleles (Mei^-^) revealed that Kar4p is required early in meiosis, before initiation of S-phase and meiotic recombination. High copy expression of the meiotic transcriptional activator *IME1* rescued the defect of these Mei- alleles. Surprisingly, Kar4p was also found to be required at a second step for the completion of meiosis and sporulation. Over-expression of *IME1* in *kar4*Δ/Δ permits pre-meiotic S-phase, but most cells remained arrested with a monopolar spindle. Analysis of the function-specific mutants revealed that roughly half became blocked after premeiotic DNA synthesis and did not sporulate (Spo^-^). Loss of Kar4p’s Spo function was suppressed by overexpression of *RIM4*, a meiotic translational regulator. Overexpression of *IME1* and *RIM4* together allowed sporulation of *kar4*Δ/Δ cells. Taken together, these data suggest that Kar4p regulates meiosis at multiple steps, presumably reflecting requirements for methylation in different stages of meiotic gene expression.

**Author Summary:** In yeast, *KAR4* is required for mating and meiosis. A genetic screen for function-specific mutations identified 25 alleles that map to different surfaces on a predicted structure of the Kar4 protein (Kar4p). The mating-specific alleles interfere with Kar4p’s ability to interact with the transcription factor Ste12p, its known partner in mating. The meiosis-specific alleles revealed an independent function: Kar4p is required for entry into meiosis and initiation of S-phase. During meiosis, Kar4p interacts with all components of the mRNA methyltransferase complex and *kar4*Δ/Δ mutants have greatly reduced levels of mRNA methylation. Thus, Kar4p is a member of the yeast methyltransferase complex. Overexpression of the meiotic transcriptional activator *IME1* rescued the meiotic entry defect but did not lead to sporulation, implying that Kar4p has more than one meiotic function. Suppression by Ime1p overexpression led to arrest after premeiotic DNA synthesis, but before sporulation. Loss of Kar4’s sporulation function can be suppressed by overexpression of a translation regulator, Rim4p. Overexpression of both *IME1* and *RIM4* allowed sporulation in *kar4*Δ/Δ cells.

## Introduction

The budding yeast, *Saccharomyces cerevisiae,* undergoes cell differentiation when it enters either the mating pathway, resulting in the fusion of two haploid cells, or meiosis, leading to sporulation and generation of haploids. Each of these pathways is mediated by specific transcription factors that activate different large-scale programs, with little overlap between their transcriptional profiles. Kar4p was originally identified as a transcription factor required for efficient yeast mating (Kurihara, Beh et al. 1994, Kurihara, Stewart et al. 1996), but it is also required for meiosis (Kurihara, Stewart et al. 1996). This raises the question of whether there are overlapping functions in the two programs.

During mating, the pheromone response triggers a mitogen-activated protein kinase cascade that activates numerous proteins, including Ste12p, the major transcription factor required for mating (Reviewed in Bardwell 2005). Ste12p turns on many genes, including a mating-specific transcript of *KAR4*, and Kar4p then works with Ste12p to activate a subset of late-acting mating genes (Kurihara, Stewart et al. 1996, Lahav, Gammie et al. 2007). The set of genes that requires Kar4p is known, but why they require Kar4p for their expression is still not well understood. Kar4p-dependent genes seem to have promoters that contain weak Ste12p- binding sites (Pheromone Response Elements, PREs) (Lahav, Gammie et al. 2007, Su, Tamarkina et al. 2010). A potential model is that Kar4p acts to stabilize Ste12p at these weak promoters. The best studied *KAR4*-dependent genes are *KAR3* and *CIK1*, which encode the two subunits of a kinesin-like motor protein responsible for nuclear congression in newly fused zygotes (Meluh and Rose 1990, Page, Satterwhite et al. 1994). When *KAR4* is absent from both mating cells, the nuclei fail to fuse (Kurihara, Stewart et al. 1996); as a result, the zygote buds off haploid daughter cells. Kar4p’s activity is Ste12-dependent, and overexpression of *STE12*, or *KAR3* together with *CIK1*, can largely suppress the *kar4*Δ mating defect (Kurihara, Stewart et al. 1996, Gammie, Stewart et al. 1999).

Similarly, meiosis is a complex process with a distinct transcriptional program; cells lacking Kar4p fail to complete the meiotic program. Entry into meiosis is regulated by a complex interplay of nutritional and cell state signals converging mainly on the promoter of *IME1*, the “early” meiotic transcription factor (Neiman 2011). Ime1p activates the expression of genes required early during meiosis for pre-meiotic DNA synthesis and the initiation of meiotic recombination. The middle meiotic transcription factor, Ndt80p, activates the expression of genes required to complete meiotic recombination and progress through the divisions and spore maturation (Neiman 2011, van Werven and Amon 2011, Winter 2012). Among the genes activated by Ime1p is *IME2,* encoding a protein kinase homologous to human CDK2 (Smith and Mitchell 1989, Kominami, Sakata et al. 1993, Guttmann-Raviv, Boger-Nadjar et al. 2001).

Ime2p both down-regulates Ime1p activity (Guttmann-Raviv, Martin et al. 2002) and phosphorylates a variety of downstream meiotic regulators (Dirick, Goetsch et al. 1998, Clifford, Marinco et al. 2004, Sedgwick, Rawluk et al. 2006, Holt, Hutti et al. 2007, Ahmed, Bungard et al. 2009, Corbi, Sunder et al. 2014). Among other functions, Ime2p is responsible for the removal of the Cdk inhibitor Sic1p, and therefore cells without *IME2* fail to initiate meiotic DNA replication (Dirick, Goetsch et al. 1998, Sedgwick, Rawluk et al. 2006).

Both the *IME1-* and *IME2*-dependent early meiotic pathways require the RNA-binding protein Rim4p (Soushko and Mitchell 2000, Deng and Saunders 2001). The early meiotic genes that are Rim4p-dependent are required for the initiation of G1 arrest, pre-meiotic DNA synthesis, and meiotic recombination (van Werven and Amon 2011). At later stages in meiosis, Rim4p blocks the translation of a set of mRNAs by sequestering them into an amyloid-like aggregate until their protein products are required (Berchowitz, Kabachinski et al. 2015, Jin, Zhang et al. 2015). Inhibition of translation is relieved by the phosphorylation of Rim4p by Ime2p (Carpenter, Bell et al. 2018) and subsequent degradation by autophagy (Wang, Zhang et al. 2020).

Although it is known that deletion of *KAR4* affects meiosis, the mechanism describing Kar4p’s function in meiosis is not known (Kurihara, Stewart et al. 1996). While *STE12* is expressed in diploids (Ellahi, Thurtle et al. 2015), it is not totally required for meiosis because cells lacking Ste12p can still form two spore asci (Marston, Tham et al. 2004). Because Kar4p’s activity in mating is Ste12p-dependent (Kurihara, Stewart et al. 1996, Gammie, Stewart et al. 1999, Lahav, Gammie et al. 2007), it is likely that Kar4p has a separate meiotic function. *KAR4* is expressed in G1 cells prior to entry into meiosis, and cells without *KAR4* fail to initiate meiosis, suggesting that it acts very early in the pathway (Kurihara, Stewart et al. 1996).

Interestingly, *KAR4* is induced again 8-12 hours into sporulation (Kurihara, Stewart et al. 1996), suggesting that it may be required at more than one stage. Clues to Kar4p’s function during meiosis can be found in its homology. Kar4p is a member of the MT-A70 family of *N*^6^- methyladenosine (m^6^A) methyltransferases (Bujnicki, Feder et al. 2002). However, Kar4p is missing several key residues important for S-adenosyl methionine binding and catalysis (Bujnicki, Feder et al. 2002, Wang, Doxtader et al. 2016, Wang, Feng et al. 2016) and is unlikely to have retained enzymatic activity. A second MT-A70 family member in yeast, *IME4*, is also required for meiosis, although unlike Kar4p, Ime4p has retained enzymatic activity (Clancy, Shambaugh et al. 2002, Agarwala, Blitzblau et al. 2012). Ime4p is part of the MIS (<u>M</u>um2p, Ime4p, and <u>S</u>lz1p) complex that methylates adenines in mRNAs during meiosis. Mutations in the *IME4* catalytic site or loss of members of the MIS complex result in reduced or delayed sporulation (Clancy, Shambaugh et al. 2002, Agarwala, Blitzblau et al. 2012). MIS complex-mediated mRNA methylation promotes expression of *IME1* by negatively regulating the expression of the transcriptional repressor, Rme1p. Methylation of *RME1* transcripts leads to their degradation, causing reduced Rme1p levels and increased expression of *IME1* (Bushkin, Pincus et al. 2019). Recently, m^6^A has been found to be widespread and dynamic during yeast meiosis, occurring in over 1000 transcripts including *IME1*, *IME2*, and *RIM4* (Bodi, Button et al. 2010, Schwartz, Agarwala et al. 2013, Scutenaire, Plassard et al. 2022, Varier, Sideri et al. 2022), which are enriched on polyribosomes (Bodi, Bottley et al. 2015).

Comparison of RNA methyltransferase homologs from different species showed that yeast *KAR4* and *IME4* diverged early in eukaryotic evolution, and both homologs are present in higher eukaryotes (Bujnicki, Feder et al. 2002). *KAR4* and *IME4* are homologous to human *METTL14* and *METTL3*, respectively. Although mRNA methylation is performed in yeast by the MIS complex, RNA methylation in humans is performed by the METTL3/ METTL14 complex together with WTAP, the human homolog of Mum2p (Agarwala, Blitzblau et al. 2012, Liu, Yue et al. 2014). Recently published crystal structures showed S-adenosyl methionine binding to METTL3, with METTL14 playing a structural role in mRNA binding (Wang, Doxtader et al. 2016, Wang, Feng et al. 2016). In yeast, Kar4p and Ime4p have been reported to physically interact, as have Kar4p and Mum2p (Uetz, Giot et al. 2000, Ito, Chiba et al. 2001, Morgan 2006), although the functional significance of the interactions is not known. Mitotic expression of the MIS complex is sufficient to induce mRNA methylation (Agarwala, Blitzblau et al. 2012), suggesting that Kar4p may not be required for methylation. However, it is important to note that Kar4p is also present in mitotic cells, thus making it possible that Kar4p was involved in the methylation they observed during mitosis in those experiments.

To better understand how Kar4p performs its functions in different developmental pathways, we conducted a genetic screen for alleles of *KAR4* that differentially affect its functions in mating and meiosis. Remarkably, Kar4p is required for at least two distinct functions in meiosis in addition to its function in mating. Kar4p’s mating function appears to be solely about its ability to interact with Ste12p, but the meiotic functions appear to be more complex. We found that *kar4*Δ/Δ mutants have serious deficiencies in mRNA m^6^A levels during meiosis and that Kar4p interacts with all members of the MIS complex. Taken together, these data imply that Kar4p is a member of the yeast methyltransferase complex. Interestingly, Kar4p, Ime4p, and Mum2p are required for another function during meiosis that appears to be independent of mRNA methylation.

## Results

### *kar4*Δ/Δ Cells Reversibly Arrest Prior to Meiotic DNA Replication

During mating, the nuclei of *kar4*Δ mutant zygotes fail to fuse, remaining separated due to the lack of induction of the kinesin motor protein Kar3p/Cik1p (Kurihara, Stewart et al. 1996). To determine where *kar4*Δ/Δ mutants are blocked in meiosis, strains expressing *GFP*-*TUB1* and *SPC42*-*mCherry* were used to observe the microtubules and spindle pole bodies by fluorescence microscopy. The progression of both wild-type and *kar4*Δ/Δ cells was visualized at 12, 24, and 48 hours after induction of meiosis (Fig 1A). Wild type cells progressed through both meiotic divisions, reaching 34% sporulation after 48 hours. In contrast, 100% of *kar4*Δ/Δ cells remain arrested with a monopolar spindle after 48 hours (Fig 1A).

**Fig 1.**
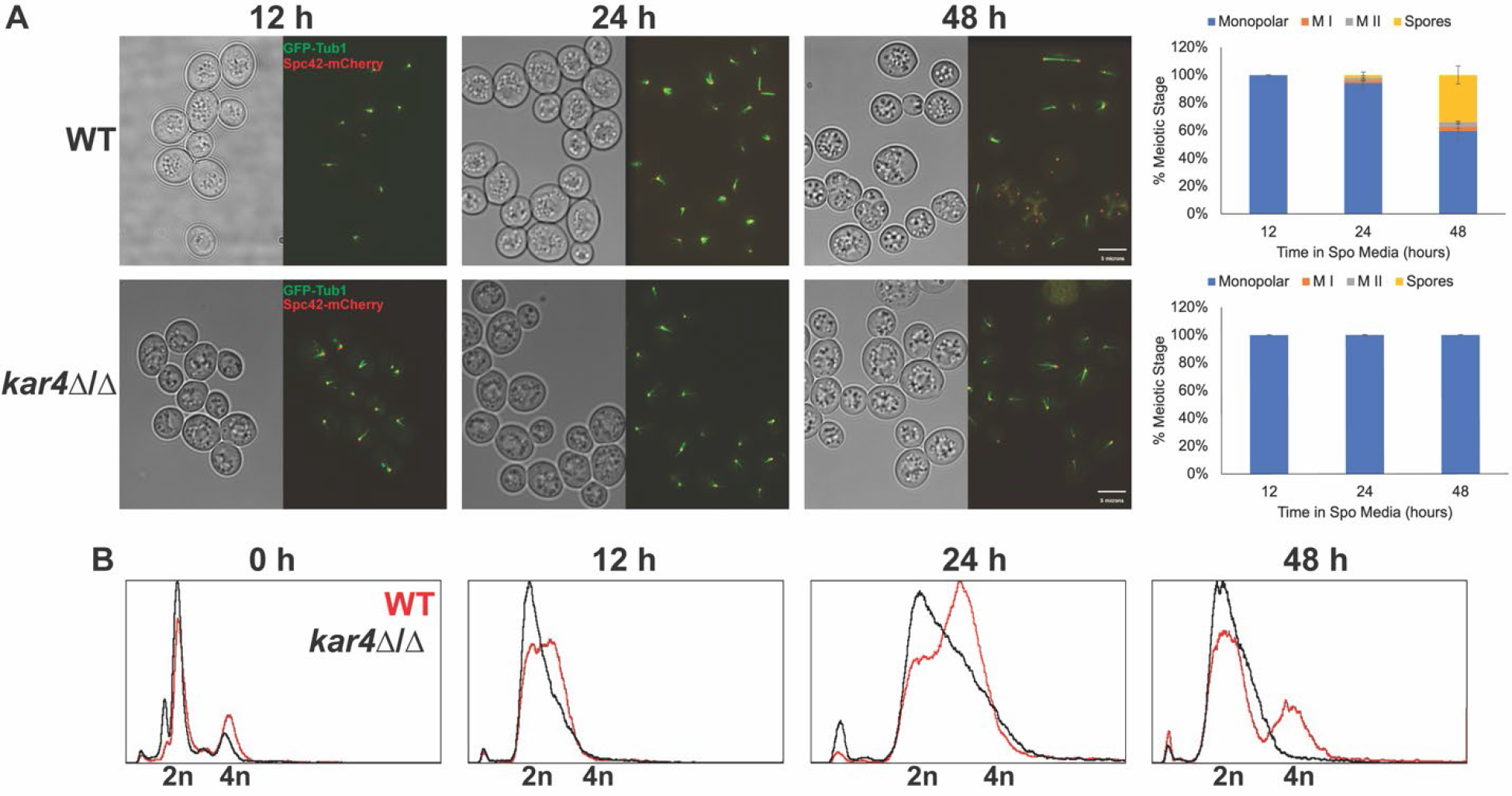
Kar4p is required early in meiosis. (A) Fluorescence microscopy of the spindle pole body (Spc42-mCherry) and microtubules (GFP- Tub1) across a time course of meiosis (12, 24, and 48 hours post transfer into sporulation media) in wild type and *kar4*Δ/Δ. Graphs are the quantification of the number of cells in different meiotic stages (Monopolar Spindle, Meiosis I, Meiosis II, and Spores). Experiments were run in three biological replicates for each strain and at least 100 cells were counted for each replicate. Error bars represent standard deviation. (B) Flow cytometry analysis of DNA content in wild type and *kar4*Δ/Δ across the same meiotic time course of the microscopy. DNA was stained with propidium iodide.

To further characterize the meiotic block in *kar4*Δ/Δ cells, we used flow cytometry to measure DNA content at 12, 24, and 48 hours. A significant number of wild-type cells entered S-phase by 12 hours; by 48 hours the cultures comprised a mixture of 2N and 4N cells. In contrast, the *kar4*Δ/Δ population comprised cells with 2N DNA content throughout the time course. We conclude that the *kar4*Δ/Δ mutants had not initiated DNA synthesis (Fig 1B), indicating that these cells were arrested very early in the meiotic program. To determine whether the arrest was reversible, the viability of the *kar4*Δ/Δ cells was assessed. The *kar4*Δ/Δ diploids remained viable after several days in sporulation media and returned to mitotic growth with near 100% efficiency.

### *KAR4* Has Different Functions in Mating and Meiosis

To determine if Kar4p has distinct functions in mating and meiosis, we designed a genetic screen for mutant alleles that might differentially affect its function in the two pathways. If Kar4p’s functions in mating and meiosis were the same, we would expect that all alleles would affect both pathways. In contrast, if Kar4p performs distinct roles in the two pathways, we would expect to isolate two classes of mutations that differentially affect function: Mat^+^ Mei^-^ and Mat^-^ Mei^+^. It is also possible that Kar4p has only one function, but that the two pathways require different levels of Kar4p activity; if so, one pathway may be more sensitive to perturbation than the other. We would then expect to isolate two classes of mutant alleles, one affecting both pathways and the other affecting only the pathway with the more stringent requirement. For example, isolation of only Mat^-^ Mei^-^ and Mat^+^ Mei^-^ alleles would suggest that *KAR4*’s meiotic function is easier to perturb than the mating function.

*KAR4* on a centromeric plasmid was mutagenized *in vitro*, and the plasmid was transformed into a *MAT***a** *kar4*Δ strain (MY 10128) (Table S1 and Table S2). To assess Kar4p’s function during mating, we used a time-limited plate mating assay (Gammie and Rose 2002), which is sensitive to defects in nuclear fusion. The transformants were replica-plated together with a *MAT*α *kar4*Δ (MY11297) (Table S1) (Tong, Evangelista et al. 2001) tester lawn onto rich media and allowed to mate for three hours before being replica-plated onto synthetic media to select for diploids. Under these conditions, the *MAT***a** *kar4*Δ control strain showed a severe mating defect (Mat^-^) compared to the wild type (Fig 2A, Fig 2B).

**Fig 2.**
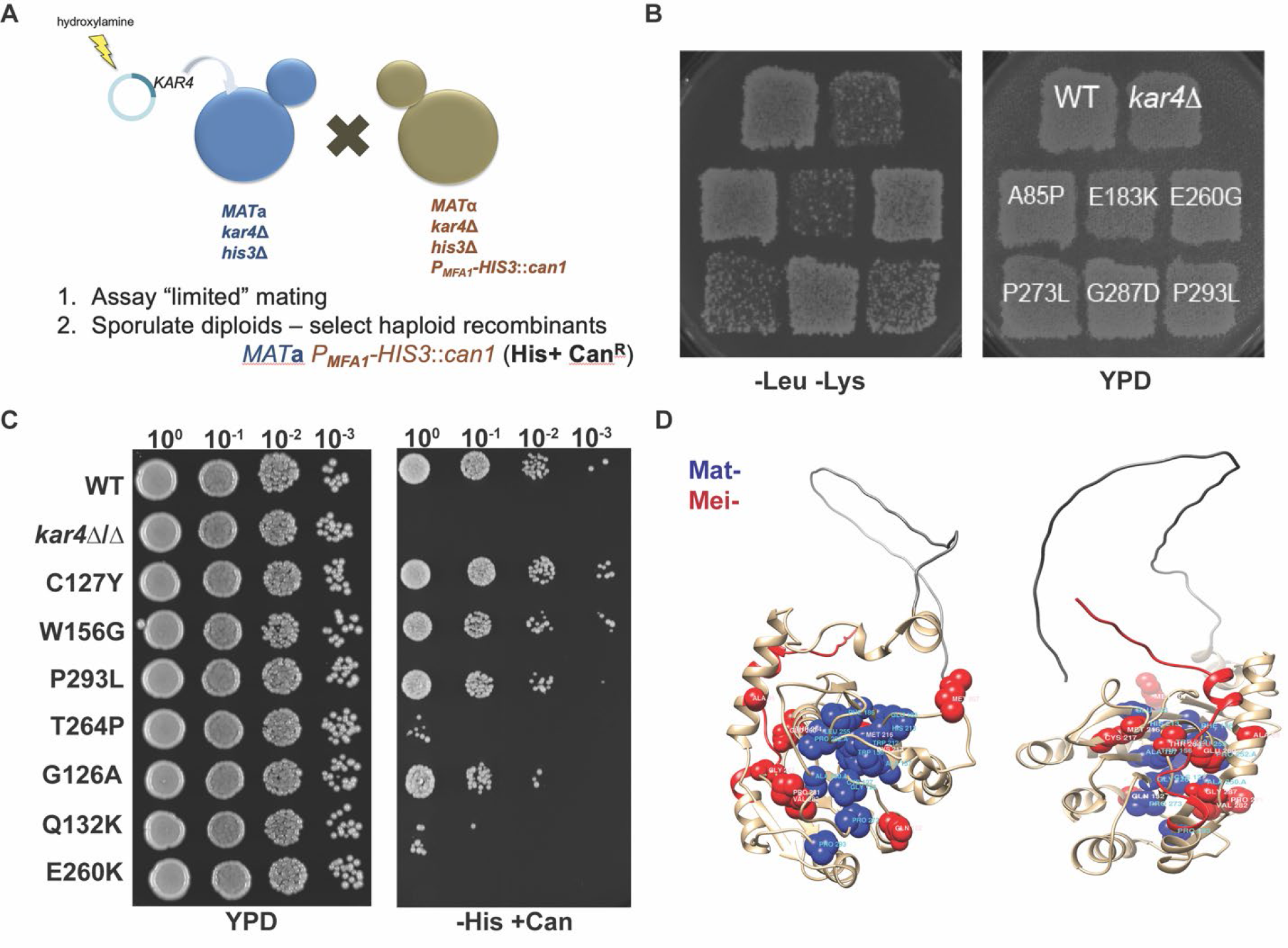
Kar4p has distinct functions in mating and meiosis. (A) Schematic of the screen used to identify separation of function mutants of *KAR4*. (B) Limited mating assay to determine the ability of the KAR4 alleles to facilitate mating. (Left) Resulting growth on media selective for diploids after a limited mating assay between a strain carrying the indicated *KAR4* allele and *kar4*Δ. (Right) Growth of strains carrying the indicated *KAR4* allele mated to *kar4*Δ on YPD (non-selective rich media) overnight. (C) Spot assay to assess the ability of the different *KAR4* alleles to initiate meiotic recombination after 24 hours of exposure to meiosis inducing conditions. (Left) Selection for successful recombination events by selecting for His+ Can^R^ recombinants. (Right) Growth of all strains on YPD. Spots are 10-fold serial dilutions of 1 OD of each sample.

**Table 1.** Characterization of the mating/meiotic defects and interactions of the separation of function alleles of *KAR4*. Mat- alleles in blue and Mei- alleles in red

| Allele | Mating | Ste12 (2-hybrid) | Meiosis | Ime4 (2-hybrid) | Mum2 (Co-IP) | % Sporulation after Ime1 OE* |
| --- | --- | --- | --- | --- | --- | --- |
| *KAR4* | + | + | ++++ | + | ++++ | 75 |
| *kar4*Δ | - | - | - | - | - | 3 |
| A157P | - | - | ++++ | ++ | ++++ | 43 |
| H213Y | - | ++ | ++++ | ++++ | ++++ | 61 |
| F186S | - | - | ++++ | +++ | + | 52 |
| E183K | - | - | ++++ | ++++ | ++++ | 52 |
| L255F | - | - | ++++ | ++++ | ++++ | 50 |
| P273L | - | - | ++++ | +++ | ++++ | 48 |
| W212R | - | ++ | ++++ | ++ | +++ | 32 |
| C127Y | - | ++ | ++++ | ++++ | ++ | 65 |
| W156G | - | - | +++ | - | ++ | 47 |
| P293L | - | - | +++ | - | ++++ | 43 |
| G126A | - | - | ++ | - | ++ | 7 |
| T264P | + | ++++ | + | ++++ | +++ | 0 |
| Q132K | + | + | + | - | ++ | 0 |
| E260K | + | + | - | ++ | + | 0 |
| G287D | + | + | - | - | - | 0 |
| E260G | + | ++++ | - | +++ | ++ | 0 |
| A310Stop | + | ++++ | - | ++ | ++ | 0 |
| ΔP281V282 | + | ++++ | - | - | - | 0 |
| M207R | + | ++++ | - | ++ | - | 0 |
| A85P | + | ++++ | - | ++++ | ++ | 0 |

\*Sporulation was measured in three biological replicates after *IME1* overexpression
\*\* “++++” ~ 100% WT, “+++” ~ 10% WT, “++” ~ 1% WT, and “+” ~ 0.1% WT

To assess Kar4p’s function in meiosis, we used a P*MFA1-HIS3* cassette integrated in place of the *CAN1* gene in the *MAT*α parent (Tong, Evangelista et al. 2001). The P*MFA1-HIS3* cassette is only expressed in cells that are phenotypically *MAT***a**. The *MAT***a** and *MAT*α parents, as well as the *MAT***a***/*α diploid, are His^-^. However, diploids competent for meiosis can produce *MAT***a** His^+^ Can^R^ haploid offspring by inducing meiotic recombination and undergoing sporulation.

Alternatively, *MAT***a** His^+^ Can^R^ diploids can be generated if cells that have undergone meiotic recombination are returned to a rich growth medium and allowed to re-enter the mitotic pathway.

After overnight incubation of the mating plate to allow multiple rounds of mating, the Mat*^-^* mutants formed a sufficient number of diploids for further analysis (Gammie and Rose 2002). The diploids were induced to enter meiosis and then spotted onto media to select for His+ Can^R^ recombinants. Under these conditions, the *KAR4*/*kar4*Δ control strains showed robust growth (Mei^+^), whereas the *kar4*Δ/*kar4*Δ diploid showed no growth (Mei^-^). Interestingly, some alleles exhibited an intermediate phenotype (Fig 2B).

A total of 25,000 transformants were screened, and three classes of mutants were identified. As expected, many mutants were defective for both mating and meiosis. As these were likely to be null mutations, they were not studied further. After rescreening and retesting, we identified 36 independent mutants representing 25 unique alleles showing separation of function (5 sites were mutated resulting in the same amino acid substitution 2 or more times). Two alleles comprised double mutants in which one or both sites were recovered as single mutations. After further analysis, 9 unique alleles were competent for mating, but not meiosis (Mat^+^ Mei^-^) and 11 alleles were defective for mating, but apparently competent for meiosis (Mat^-^ Mei^+^). Because we identified all three possible classes of mutant alleles (null mutations and both types of separation of function alleles), we conclude that Kar4p has distinct, separable functions in mating and meiosis (Table 1).

To identify the causal mutations in *KAR4*, the plasmids were recovered and sequenced. The alleles were interspersed throughout the primary sequence of *KAR4*, suggesting that Kar4p is not organized into distinct functional domains (Table 1, S Fig 1). To determine if the function-specific alleles map to different surfaces of the protein, we took advantage of the recent advances in protein folding algorithms to generate AlphaFold (Jumper, Evans et al. 2021) predicted structure of Kar4p and mapped the alleles to that structure (Fig 2C). The two classes of alleles clustered into distinct regions of the predicted protein. The Mat- alleles (blue) cluster tightly within one region, whereas the Mei- alleles (red) are found broadly distributed over the opposite surface. The finding that distinct regions of the predicted protein are affected by the two classes of mutations suggests that these regions mediate different functions in mating and meiosis.

### Kar4p Engages in a Function Specific Interaction with Ste12p

One possible model for how the mutations differentially impact Kar4p’s functions is that they block the ability of Kar4p to interact with different protein partners. Kar4p’s activity during mating is dependent on Ste12, over-expression of Ste12p suppresses the loss of Kar4p (Kurihara, Stewart et al. 1996), and Kar4p interacts with Ste12p during the pheromone response (Lahav, Gammie et al. 2007). Therefore, it is likely that the *kar4* Mat^-^ alleles interfere with mating by disrupting the Kar4p-Ste12p interaction. To test the ability of the *KAR4* alleles to interact with Ste12p, we leveraged Ste12p’s function as a transcriptional activator to perform a one-hybrid assay between the Kar4p mutants and Ste12p. We moved the alleles onto a construct of *KAR4* fused to the Gal4p DNA-binding domain (*KAR4*-GBD) previously used to show that Kar4p’s mating activity is Ste12p-dependent (Lahav, Gammie et al. 2007). In the presence of mating pheromone, the Kar4 hybrid protein recruits Ste12p to drive transcription of a *HIS3* reporter gene, allowing cells to grow in the absence of histidine. In strains carrying alleles that disrupt the Kar4p-Ste12p interaction, the reporter will not be transcribed. Of the 20 mutants tested, 8 of the 11 Mat- alleles caused reduced interaction with Ste12p. All 9 of the Mat+ alleles allowed transcription of the reporter gene (Fig 3A, Table 1). Interestingly, three alleles were defective for mating but still showed some interaction with Ste12p by this assay. The presence of these alleles suggests that there may be some threshold of Kar4p function that is required for mating or that Kar4p may do more than just stabilize Ste12p at weak promoters during mating. We conclude that most alleles that interfere with mating function do so by disrupting the Kar4p-Ste12p interaction.

**Fig 3.**
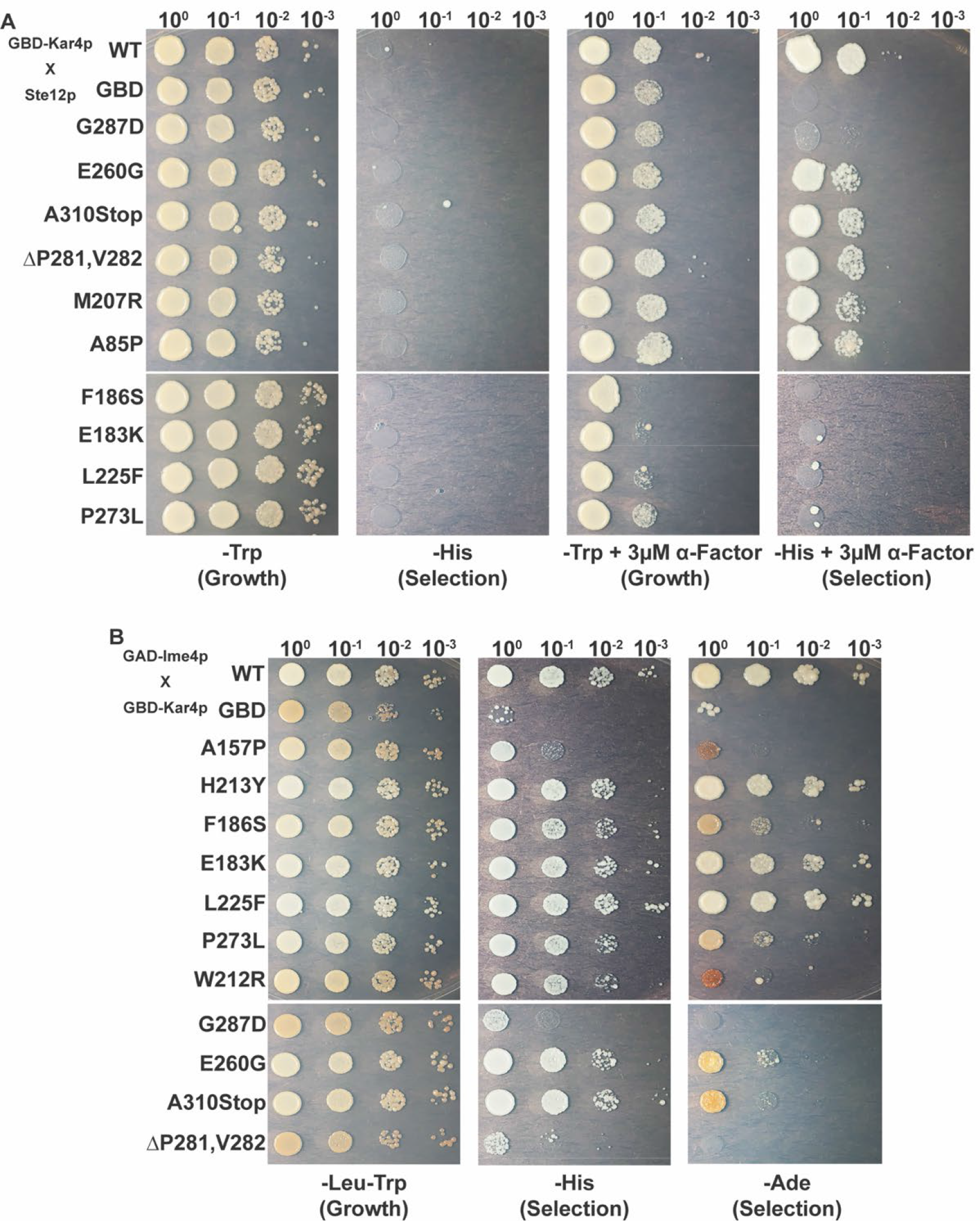
Kar4p engages in a function specific interaction with Ste12p and Ime4p. (A) A one-hybrid assay to determine the ability of the different Kar4p alleles fused to the Gal Binding Domain to interact with Ste12p upon exposure to alpha-factor. (Left) Growth on non-selective media both with and without 3 µM alpha-factor. (Right) Growth on SC-His media to select for alleles that can maintain an interaction with Ste12p and drive expression of the *HIS3* reporter gene. (B) A two-hybrid assay was used to determine the ability of the different Kar4p alleles to interact with Ime4p fused to the Gal Activating Domain after 24 hours of exposure to meiosis inducing conditions. (Left) Growth on non-selective media. (Right) Growth on both SC-HIS and SC-ADE to select for the alleles that can maintain an interaction with Ime4p and drive expression of the two reporter genes *HIS3* and *ADE2*. Spots are 10-fold serial dilutions of 1 OD of each sample.

### Kar4p’s Interaction with Ime4p is Correlated with the Mei Function

It seemed likely that Kar4p’s role in meiosis might also involve a function specific binding partner. Ime4p, the paralog of Kar4p, was a good candidate for several reasons: (1) the orthologs of Ime4p and Kar4p in most eukaryotes have been shown to form a heterodimer that functions to methylate mRNA, (2) both *ime4*Δ/Δ and *kar4*Δ/Δ arrest at similar points in meiosis, and (3) a physical interaction between Kar4p and Ime4p was previously detected by high-throughput yeast two-hybrid assays (Ito, Chiba et al. 2001). To confirm the interaction, we created a fusion of *IME4* to the Gal4p activating domain (*IME4*-GAD) for use in the yeast two-hybrid system with the *KAR4*-GBD fusion described above. Our findings agree with the previous report that Kar4p and Ime4p physically interact (Fig 3B). We also detected the interaction when the fusions were swapped (*KAR4*-GAD with *IME4*-GBD) (data not shown).

To identify residues on Kar4p important for the interaction with Ime4p, we tested the alleles in the two-hybrid assay. All 8 alleles that are proficient for wild type levels of meiotic function were able to drive varying levels of expression of the reporter gene. Of the four alleles that had reduced meiotic function, three were not able to express the reporter gene. Of the 12 alleles that showed either reduced or total loss of meiotic function, 6 were not able to interact with Ime4p and the other 6 had varying levels of expression of the reporter gene (Fig 3B, Table 1). One interpretation of these data is that the alleles causing a defect in meiotic function, but not in the interaction with Ime4p, may nevertheless impact the function of Kar4p in meiosis. We conclude that Kar4p’s role in meiosis is correlated with its interaction with Ime4p, implying that Kar4p may also have a role in mRNA methylation.

### Kar4p is Required for Efficient mRNA Methylation During Meiosis

To address whether Kar4p is required for mRNA methylation during meiosis, we measured m^6^A levels in mRNAs from sporulating cells. To increase the efficiency and synchrony of meiosis, we moved the *kar4*Δ/Δ, *ime4*Δ/Δ, and *slz1*Δ/Δ into the SK1 background.

Cells were harvested after four hours in sporulation-inducing conditions, which we determined was when m^6^A levels peaked during meiosis in the strains we used for these experiments (data not shown). In agreement with previous findings, we were able to detect m^6^A levels at 25% of wild type in the *slz1*Δ/Δ strain. (Fig 4A). Levels of m^6^A were only 17% of wild type in *kar4*Δ/Δ indicating that Kar4p is required for wild type levels of mRNA methylation during meiosis.

**Fig 4.**
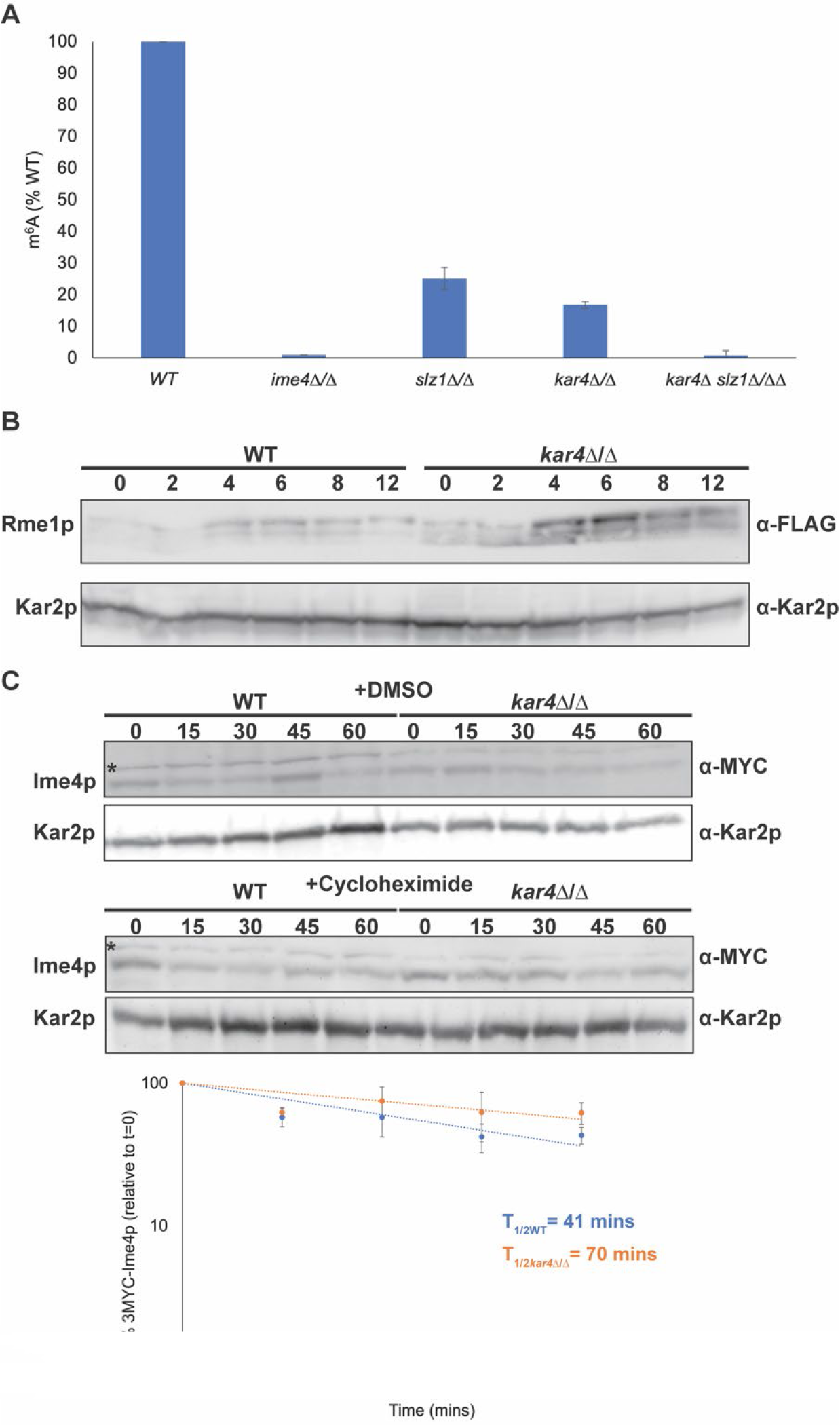
Kar4p is required for mRNA m^6^A methylation. (A) mRNA m^6^A levels measured using an ELISA like assay from EpiGenTek. The indicated mutations were made in the SK1 strain background and samples were harvested after four hours of exposure to meiosis inducing conditions. Experiments were run in three biological replicates for each strain and error bars represent standard deviation. (B) Western Blot of 3xFLAG-Rme1p in wild type and *kar4*Δ/Δ across a time course of meiosis. Kar2p is used as a loading control. (C) Western blot of 3xMYC-Ime4p after four hours in meiosis inducing media with either 100 µM cycloheximide or an equivalent amount of DMSO in both wild type and *kar4*Δ/Δ. (Top) 3xMYC-Ime4p levels with DMSO. (Middle) 3xMYC-Ime4p levels with cycloheximide. (Bottom) Quantification of three biological replicates of the cycloheximide chase experiment. The strains used are in the SK1 background. Kar2p is used as a loading control. “*” indicates a non-specific band.

Interestingly, in *kar4*Δ/Δ *slz1*Δ/Δ double mutants we were unable to detect methylated mRNA. Slz1p is important for localization of the MIS complex to the nucleus (Schwartz, Agarwala et al. 2013), but some nuclear localization persisted in *slz1*Δ/Δ. Given that Kar4p localizes to the nucleus during mating, it is tempting to speculate that nuclear localization of the other members of the MIS complex persisted in the *slz1*Δ/Δ mutant in meiosis due to Kar4p.

Meiotic mRNA methylation reduces the levels of the transcriptional repressor Rme1p. In mutants lacking Ime4p (or catalytic mutants of Ime4p), Rme1p levels are increased causing lower levels of *IME1* transcription (Bushkin, Pincus et al. 2019). Consistent with a role for Kar4p in mRNA methylation, the levels of Rme1p tagged with three FLAG repeats remained high in *kar4*Δ/Δ across meiosis, elevated at least 2-fold for the duration of the time course (Fig 4B).

Kar4p may be required for mRNA methylation either because it is required for the activity of the complex, or by stabilizing Ime4p. By using cycloheximide to block new protein synthesis during meiosis, we found that Ime4p does not turn over faster in the absence of Kar4p (Fig 4C). Indeed, Ime4p levels were slightly higher in *kar4*Δ/Δ compared to wild type, which could be due to the delay in sporulation in those mutants. We conclude that Kar4p is required for the activity of the methyltransferase complex and not for stabilizing Ime4p.

### Kar4p Interacts with the MIS Complex Proteins Mum2p and Slz1p

If Kar4p is a member of the methyltransferase complex, it should also interact with the other known components, Mum2p and Slz1p. A high throughput study using the two-hybrid method suggested an interaction between Kar4p and Mum2p (Uetz, Giot et al. 2000). To confirm this, we used co-immunoprecipitation of Kar4p fused to the 3xHA epitope and Mum2p fused to the 4xMYC epitope. We were able to confirm an interaction between Kar4p and Mum2p (Fig 5A). In addition, we used the alleles of *KAR4* and co-immunoprecipitations to ask whether Kar4p and Mum2p engage in a function specific interaction during meiosis. Interaction of the Kar4p mutant proteins with Mum2p was highly correlated with the interaction of Kar4p with Ime4p.

**Fig 5.**
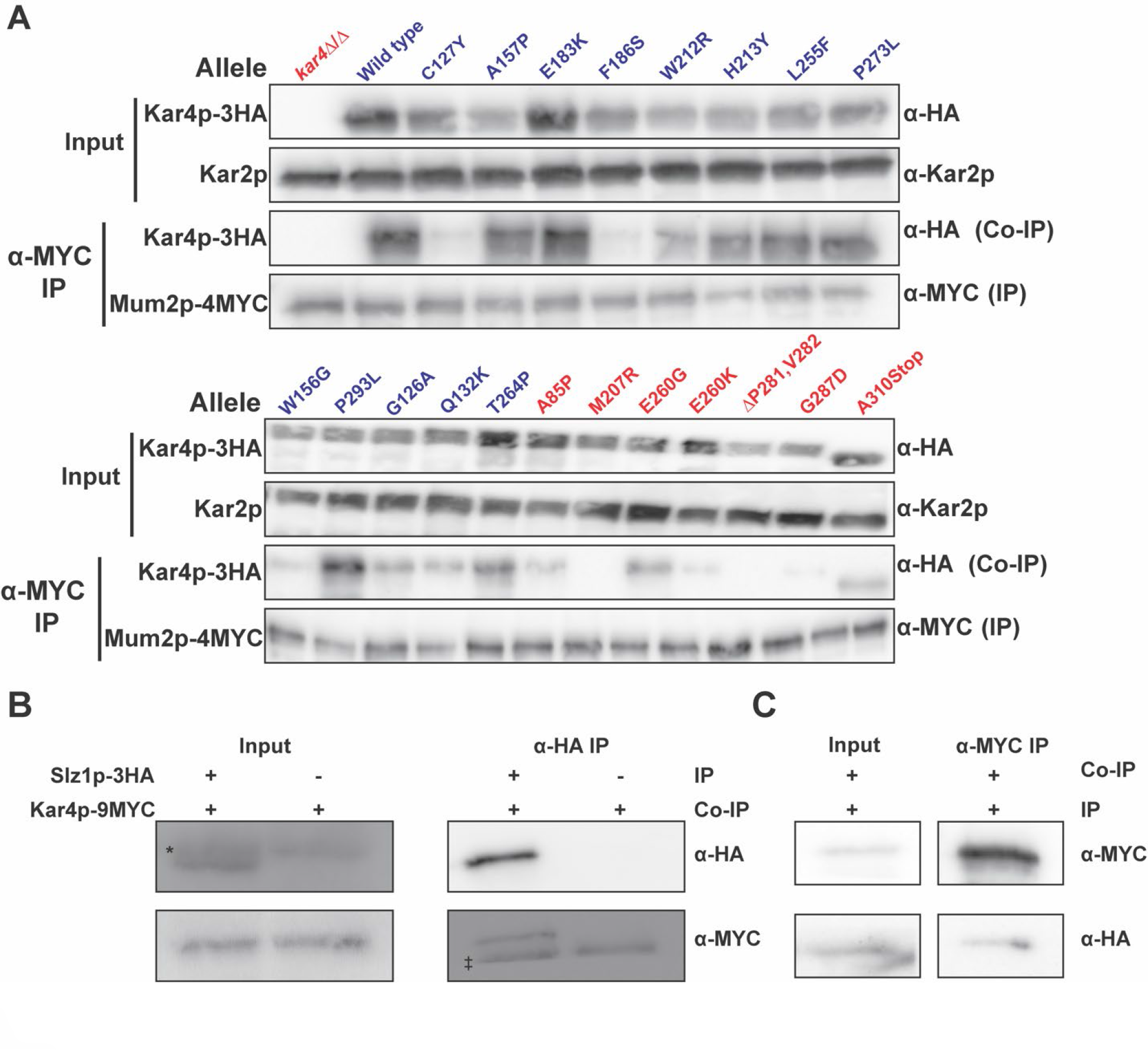
Kar4p interacts with the MIS complex components Mum2p and Slz1p. (A) Western blots of total protein and co-IPs between Mum2p-4MYC and the alleles of Kar4p- 3HA. (Top) Total protein samples from the extracts that were used for the Co-IPs. Kar2p is used as a loading control. Alleles proficient for Kar4p’s meiotic function (Mei^+^) are in blue and alleles in red are not (Mei^-^). (Bottom) Co-IPs where Mum2p-4MYC was purified and the co- purification of Kar4p-3HA was assayed. (B) Western blots of total protein and Co-IPs between Slz1p-3HA and Kar4p-9MYC. (Left) Total protein samples from the extracts that were used for the co-IPs. “*” indicates a non-specific band. (Right) Co-IPs where Slz1p-3HA was purified and the co-purification of Kar4p-13MYC was assayed. “‡” indicates the heavy chain of IgG from the anti-HA magnetic beads used for the Co-IP. (C) Western blots of total protein and Co-IPs between Kar4p-9MYC and Slz1p-3HA. (Left) Total protein samples from the extracts that were used for the Co-IPs. (Right) Co-IPs where Kar4p-9MYC was purified and the co-purification of Slz1-3HA was assayed.

Only 4 out of the 20 mutant Kar4p proteins tested showed an interaction with Mum2p that was different from that observed with Ime4p (Fig 5A, Table 1). Three mutant proteins (P293L, G126A, and Q132K) interacted with Mum2p and retained some meiotic functionality, but were strongly defective for interaction with Ime4p, by the two-hybrid assay. One possibility is that there is still an interaction between the Kar4p and Ime4p that is facilitated by the interaction with Mum2p. This may explain why these alleles retain some meiotic function. The fourth anomalous mutant protein, M207R, maintained an interaction with Ime4p, but was defective for both meiotic function and interaction with Mum2p. This suggests that interaction with Ime4p alone is not sufficient for facilitating Kar4p’s meiotic functions. Taken together, Kar4p’s meiotic function appears dependent upon interaction with two key members of the MIS complex, Ime4p and Mum2p.

To determine if Kar4p also interacts with Slz1p we generated strains carrying Slz1p fused to the 3xHA epitope and Kar4p fused to the 13xMYC epitope and performed co-immunoprecipitation assays during meiosis. We detected interaction between the two proteins either by precipitating Slz1p or Kar4p (Fig 5B and Fig 5C). We did not examine whether the interaction is affected by the function specific *KAR4* alleles. Thus, Kar4p interacts with all members of the MIS complex and is required for mRNA methylation. These findings point to Kar4p being a member of the mRNA methyltransferase complex.

### Kar4p’s Meiotic Function Is Partially Suppressed by Overexpression of *IME1*

To gain insight into Kar4p’s meiotic function, we performed a high-copy suppressor screen. The strain used in the initial mutant screen (MY10128) was transformed with a YEp24 2µ plasmid-based genomic library (Carlson and Botstein 1982) and the transformants were mated to the *kar4*Δ *MAT*α *can1*::PMFA1*HIS3* strain used to discover the Mei- alleles. Colonies that could now undergo meiotic recombination were identified after induction of sporulation by growth on appropriate selective media (SC-histidine, plus canavanine). The *kar4*Δ/Δ recombination defect was suppressed by 2µ plasmids carrying chromosomal fragments containing *KAR4* (6x)*, IME1* (12x), and *IME2* (1x). To confirm the identity of the suppressing genes, *IME1* and *IME2* were subcloned onto 2µ vectors and retested (Fig 6A). Suppression by *IME1* was not as strong as *KAR4*, suggesting that Kar4p may have additional functions other than acting through Ime1p. Suppression by *IME2* was much weaker than *IME1* (Fig 6A).

**Fig 6.**
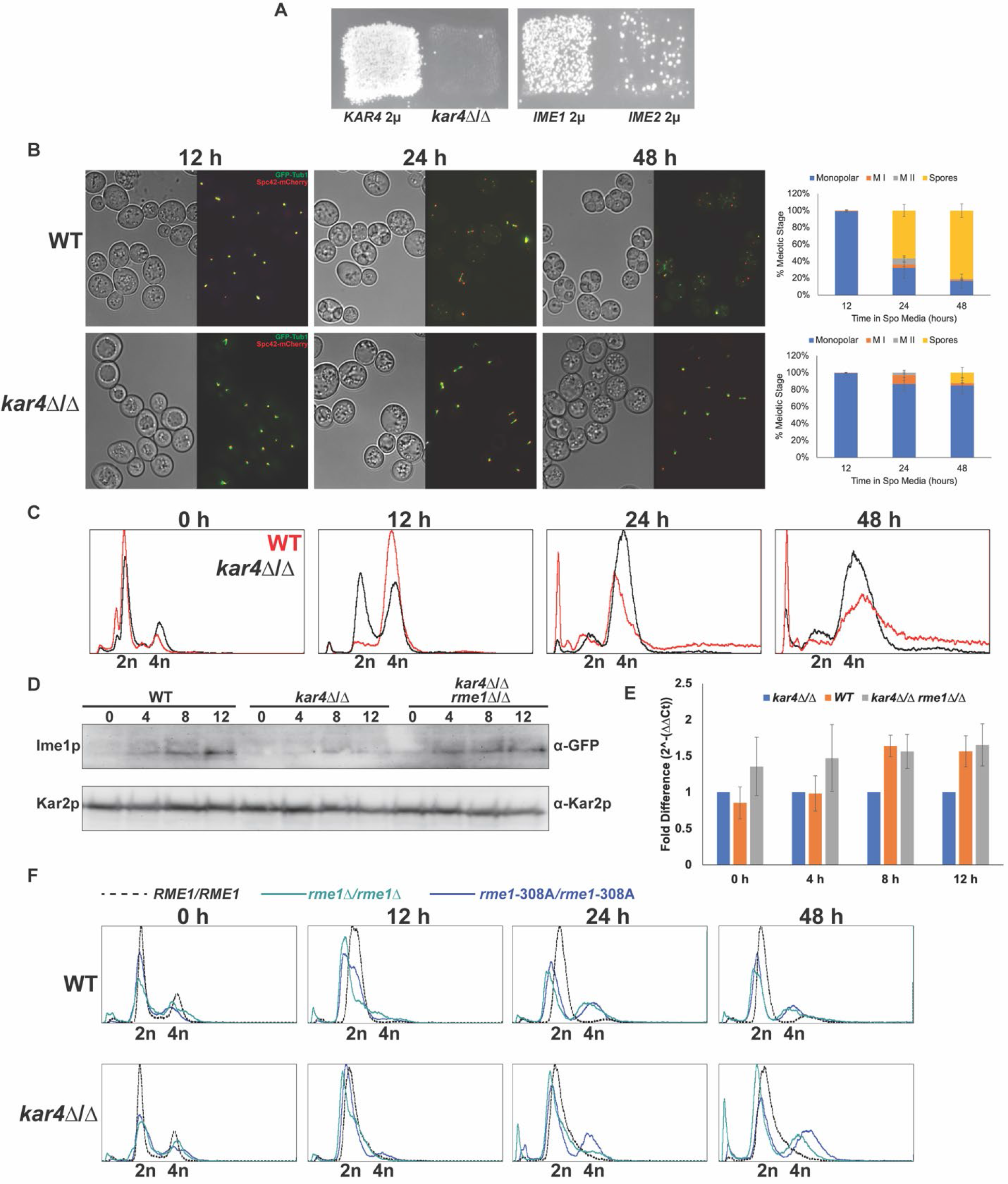
*IME1* overexpression partially suppresses the *kar4*Δ/Δ meiotic defect. (A) Growth of *kar4*Δ/Δ cells on SC-HIS+canavanine to screen for recombination after exposure to meiosis-inducing conditions carrying either *KAR4*, *IME1*, or *IME2* cloned onto high-copy number 2µ plasmids. (B) Fluorescence microscopy of the spindle pole body (Spc42-mcherry) and microtubules (GFP-Tub1) across a time course of meiosis (12, 24, and 48 hours post movement into sporulation media) in wild type and *kar4*Δ/Δ with *IME1* overexpressed from an estradiol-inducible promoter. Graphs are the quantification of the number of cells in different meiotic stages (Monopolar Spindle, Meiosis I, Meiosis II, and Spores). Experiments were run in three biological replicates for each strain and at least 100 cells were counted for each replicate. 1 µM of estradiol was used to induce expression. Error bars represent standard deviation. (C) Flow cytometry analysis of DNA content in wild type and *kar4*Δ/Δ with *IME1* overexpressed across the same meiotic time course of the microscopy. DNA was stained with propidium iodide. (D) Western blot showing GFP-Ime1p levels across a time course of meiosis in wild type, *kar4*Δ/Δ, and *kar4*Δ*rme1*Δ/ *kar4*Δ*rme1*Δ. Kar2p is used as a loading control. (E) qPCR measuring *IME1* transcript levels in wild type, *kar4*Δ/Δ, and *kar4*Δ*rme1*Δ/ *kar4*Δ*rme1*Δ. Data were normalized to levels of *PGK1* expression. (F) Flow cytometry analysis of DNA content in wild type and *kar4*Δ/Δ carrying either the wild type allele of *RME1*, the hypomorphic -308A allele of *RME1*, or a deletion of *RME1*. DNA was stained with propidium iodide.

To further investigate *IME1*’s suppression of the *kar4*Δ/Δ meiotic phenotype, we used a non-native, estradiol-inducible promoter to overexpress *IME1* in cells containing GFP-*TUB1* and *SPC42*-mCherry. In this system (McIsaac, Gibney et al. 2014), the promoter is recognized by an artificial transcription factor with four zinc fingers (Z4EV), effectively eliminating any off-target effects. Addition of estradiol triggered rapid sporulation in wild-type diploid cells (*P*Z4EV-*IME1*) resulting in 57% sporulation after 24 hours and 81% sporulation by 2 days. In contrast, induction of PZ4EV *-IME1* in *kar4*Δ/Δ cells resulted in no sporulation after 24 hours, only 7% sporulation after 48 hours, and reached a peak of 12% sporulation after 2 days. Most cells remained arrested with a monopolar spindle (Fig 6B), however almost all cells had completed premeiotic S-phase (Fig 6C). After induction of PZ4EV-*IME1, KAR4* cells retained 94% viability. In contrast, the PZ4EV-*IME1-*induced *kar4*Δ/Δ strains retained only 27% viability (S Fig 2).

We conclude that *IME1* overexpression allows premeiotic S-phase, consistent with the appearance of meiotic recombinants. However, only a small fraction of cells complete sporulation in the *kar4*Δ/Δ cells, indicating that there are remaining defects in meiosis that are not suppressed by *IME1* overexpression. Given Kar4p’s role in RNA methylation and the role of RNA methylation in meiotic entry, we hypothesize that the requirement for Kar4p in meiotic DNA synthesis and recombination is through the expression of *IME1*. In support of this hypothesis, Ime1p levels were reduced at least 2-fold in *kar4*Δ/Δ compared to wild type over the first 12 hours of meiosis (Fig 6D). A much less severe impact was observed for the level of *IME1* transcript (Fig 6E). Interestingly, there was no difference in the level of *IME1* transcript at 4 hours between wild type and *kar4*Δ/Δ, although there was a significant defect in the level of Ime1p protein. Recent work confirmed that *IME1* transcripts are methylated (Scutenaire, Plassard et al. 2022, Varier, Sideri et al. 2022), suggesting that the loss of methylation in *kar4*Δ/Δ may impact the translatability of *IME1* mRNA. Together, these data suggest that Kar4p is required for the normal induction of *IME1* expression.

### Kar4p’s Meiotic Function is Partially Suppressed by Loss of Rme1p

Recent work showed that RNA methylation acts upstream of *IME1* expression through regulating the levels of Rme1p (Bushkin, Pincus et al. 2019). RNA methylation acts to destabilize *RME1* transcript and thereby reduce the levels of Rme1p. Reducing Rme1p levels either by hypomorphic allele (*RME1-308A*) found in the SK1 strain background or by deleting *RME1* permits meiotic DNA replication in *ime4*Δ/Δ, but does not permit sporulation (Bushkin, Pincus et al. 2019). Given Kar4p’s role in RNA methylation and its impact on Ime1p levels, we sought to determine if loss of Rme1p could also permit meiotic DNA replication in *kar4*Δ/Δ. Wild type strains harboring either the hypomorphic allele or the deletion underwent meiotic replication faster than strains carrying the wild type *RME1* allele (Fig 6F). In *kar4*Δ/Δ with these mutations, cells also underwent meiotic replication, although not to the same extent that occurs upon overexpression of *IME1* (Fig 6F). Like *IME1* overexpression, the *RME1* mutations did not permit spore formation in *kar4*Δ/Δ (S Fig 3). A double mutant lacking both Kar4p and Rme1p expressed wild type levels of both *IME1* transcript and protein (Fig 6D, Fig 6E). Again, we note that rather small changes in mRNA levels in the double mutant (compared to *kar4*Δ/Δ alone) resulted in much larger increases in protein level. Possibly *IME1* is also regulated at the translational level, by a mechanism other than mRNA methylation, or the double mutant may have increased expression of *IME4,* and increased levels of mRNA methylation compared to *kar4*Δ/Δ. These findings, in conjunction with the ability of *IME1* overexpression to rescue the early *kar4*Δ/Δ defect, further support the conclusion that Kar4p acts upstream of *IME1* expression. Increasing Ime1p through overexpression of *IME1* or by removing a negative regulator can bypass the early requirement for Kar4p and mRNA methylation.

### Kar4p has Two Distinct Meiotic Functions

We next tested whether overexpression of *IME1* would rescue the separation of function alleles described above. Plasmids bearing the alleles were transformed into a diploid *kar4*Δ/Δ strain carrying the PZ4EV-*IME1* construct. Transformant strains were incubated in sporulation media, induced with estradiol, and examined for spore formation after 48 hours. As expected, many of the Mei^-^ mutants were unable to form spores, like *kar4*Δ/Δ. Remarkably, two of the mutants (W156G and P293L) that show reduced meiotic function were able to form spores at wild type levels after overexpression of *IME1*. However, the three alleles that had even greater reduction in meiotic function (G126A, T264P, and Q132K) were not able to sporulate at wild type levels after *IME1* overexpression (Table 1). Given the high degree of correlation between the alleles that are both Mei^+^ and Spo^+^, it is difficult to discern whether the Spo function is a distinct meiotic function of Kar4p or simply requires higher levels of methyltransferase activity.

To elucidate whether the Mei and Spo functions are distinct functions of Kar4p, we examined the ability of *IME1* overexpression to suppress the meiotic defects caused by deletions of other members of the methyltransferase complex. If the meiotic defects can be rescued by *IME1* overexpression alone, this would suggest that the Spo function may be a distinct meiotic function. In our strains, deletions of *KAR4, IME4,* and *MUM2* completely fail to sporulate; the *slz1*Δ/Δ had low levels of sporulation (S Fig 4). Overexpression of *IME1* did not rescue sporulation in *ime4*Δ/Δ or *mum2*Δ/Δ mutants (Fig 7, S Fig 4). However, the slz*1*Δ/Δ mutant was suppressed by overexpressing *IME1*. A catalytic mutant of Ime4p (*ime4-D348A, W351A* referred to as *ime4-cat* from this point forward) lacks Ime4p methyltransferase activity and has been shown to block meiosis (Clancy, Shambaugh et al. 2002). Interestingly, the catalytic mutant was rescued by overexpression of *IME1* alone, suggesting that the lack of methylation is largely manifest as a defect in Ime1p function. This further suggests that Kar4p’s Spo function does not involve RNA methylation and represents a distinct meiotic function of Kar4p (Fig 7, S Fig 4). By extension, this finding implies that Slz1p’s function in meiosis is only through its role in mRNA methylation.

**Fig 7.**
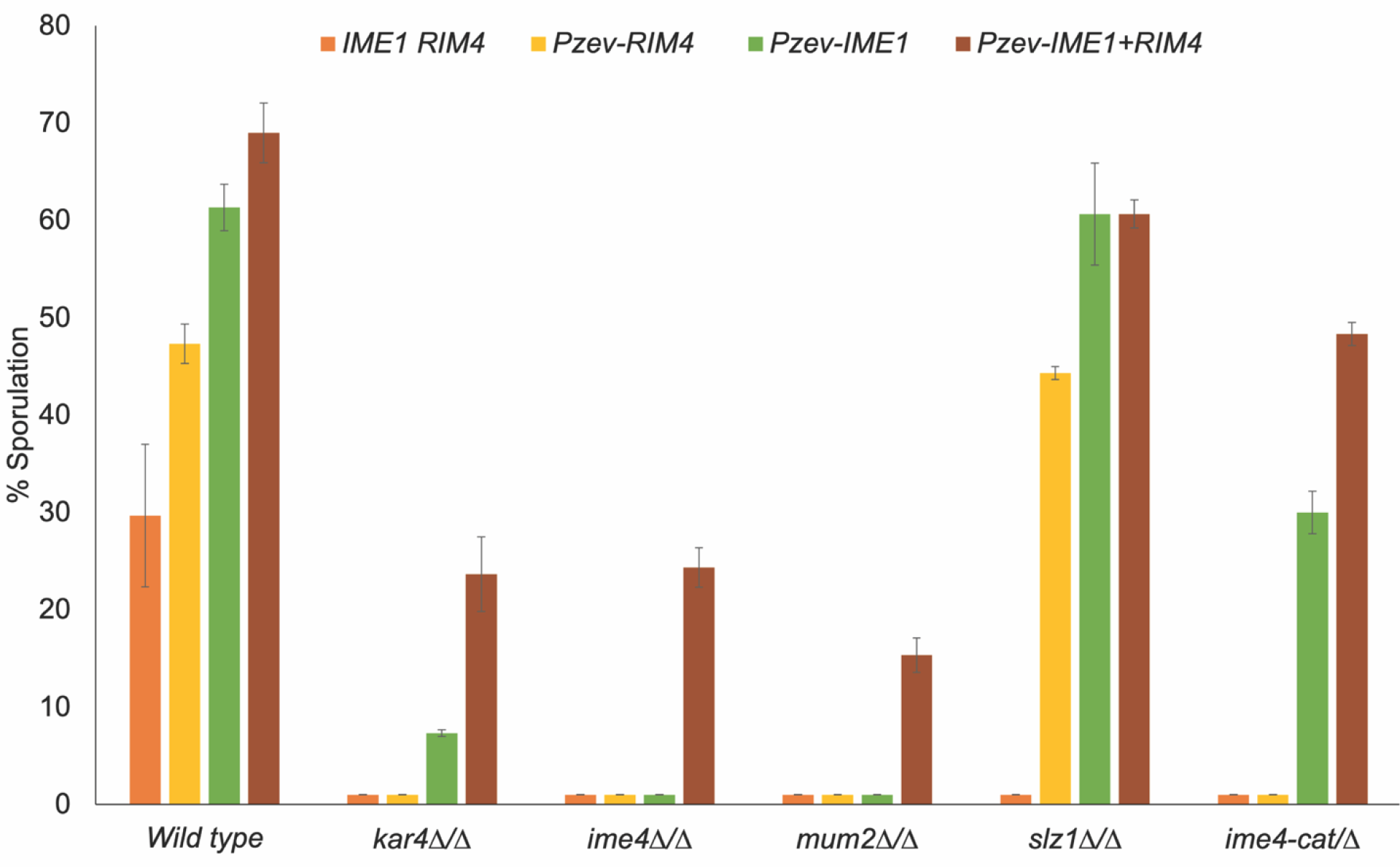
Co-overexpression of *IME1* and *RIM4* permits sporulation in *kar4*Δ/Δ. Sporulation of wild type, *kar4*Δ/Δ, *mum2*Δ/Δ, *ime4*Δ/Δ, *slz1*Δ/Δ, and *ime4-cat*/*ime4*Δ with either endogenous expression of *IME1* and *RIM4*, overexpression of *RIM4*, overexpression of *IME1*, or overexpression of both *IME1* and *RIM4*. 1 µM of estradiol was used to induce expression. All dyads, triads, and tetrads were counted across three biological replicates for each strain. At least 100 cells were counted after 48 hours post addition of estradiol. Error bars represent standard error.

Taken together, these observations indicate that Kar4p is required at more than one step during meiosis. An early step that is upstream of meiotic DNA synthesis and recombination (Kar4p’s Mei function) appears to be associated with its role in RNA methylation. A later step acts either during or downstream of recombination but upstream of commitment to meiosis, given the ability of *kar4*Δ/Δ mutants to resume mitotic growth after *IME1* overexpression. The later step (Kar4p’s Spo function) appears to be a distinct function of Kar4p and other members of the methyltransferase complex, other than Slz1p, that does not involve RNA methylation. Given the role of Slz1p in localizing the complex to the nucleus, the non-catalytic function may be specific to the cytoplasm.

### Overexpression of *RIM4* with *IME1* leads to spore formation in *kar4*Δ/Δ

To better understand Kar4p’s “Spo” function, we designed an unbiased high-copy suppressor screen to look for genes that could rescue the sporulation defect of PZ4EV-*IME1* induced *kar4*Δ/Δ diploids. We transformed the same 2μ library used previously into the PZ4EV- *IME1 kar4*Δ/Δ diploid strain. Transformants were washed off plates and induced to sporulate by addition of β-estradiol. After 3 days, the cultures were digested with zymolyase, which kills diploids that do not complete meiosis, but spores are resistant to this treatment. Survivors were plated on media selecting for the suppressing plasmid. Plasmids were extracted from haploid colonies and their inserts identified by DNA sequencing. From 13 independent transformant colonies, we recovered 5 different *KAR4*-containing genomic fragments. From 3 other independent transformants, we recovered the same 7.5 kb fragment from chromosome VIII. Sequence analysis determined that it contained the sequences for *RIM4, SNF6, YHL026C*, and a portion of the *RIM101* gene. Subcloning of the genomic fragments onto new 2μ plasmids identified *RIM4* as the suppressor gene. For all further experiments, we put *RIM4* and *IME1* under the control of an estradiol-inducible promoter (PZ3EV). In confirmation PZ3EV-*IME1*, PZ3EV- *RIM4* induced *kar4*Δ/Δ strains were able to form spores, while the uninduced strains did not (Fig 7, S Fig 4). Overexpression of *RIM4* alone was able to permit sporulation in *slz1*Δ/Δ, but not any of the other mutants (Fig 7, S Fig 4).

Given the ability of *IME1* and *RIM4* co-overexpression to rescue the *kar4*Δ/Δ defect, we asked if it could also rescue the *ime4*Δ/Δ and *mum2*Δ/Δ meiotic defects. For both mutants, the co-overexpression of *IME1* and *RIM4* was able to permit sporulation at levels comparable to those seen in *kar4*Δ/Δ (Fig 7, S Fig 4). Taken together, these data suggest that Kar4p, Ime4p, and Mum2p may all participate in a second RNA methylation-independent meiotic function. The requirement for *RIM4* overexpression suggests that this function may involve translational regulation of meiotic transcripts.

## Discussion

We took advantage of the power of yeast genetics to understand how *KAR4* is integrated into multiple differentiation pathways in yeast. The genetic screen for separation-of-function mutants identified multiple alleles that specifically disrupt either Kar4p’s mating or meiotic function. These alleles are interspersed on Kar4p’s primary sequence, suggesting that it does not have separate functional domains. However, the three-dimensional model threaded using the AlphaFold predicted structure of Kar4p (Jumper, Evans et al. 2021) shows that these alleles cluster in separate regions of the protein.

The mechanism of Kar4p’s mating function has been relatively well-characterized. It is known to interact with Ste12p to activate a subset of pheromone-induced genes, specifically those with “weak” PREs, which tend to be expressed later in mating and at higher pheromone concentration. Although Kar4p does not bind DNA directly, electrophoretic-mobility shift assays suggest that it facilitates the binding of Ste12p to Kar4p-dependent promoters. Evidence for direct interaction comes from the ability of Kar4p to recruit Ste12p and activate transcription in a one-hybrid transcription assay (Lahav, Gammie et al. 2007). All Mat^+^ mutants interact with Ste12p by this assay, and all but three of the Mat- mutants were fully defective for interaction.

Thus, we conclude that the Mat^-^ mutations are specifically defective for mating because they fail to interact with Ste12p and that Mat^-^ alleles likely define a region on Kar4p that is required for binding Ste12p. Furthermore, these data in combination with the fact that loss of Ste12p does not fully block meiosis indicate that the Kar4p-Ste12p interaction is not required for meiosis.

In meiosis, Kar4p may have two different functions. It is first required prior to pre-meiotic S-phase and meiotic recombination. Kar4p is required for efficient mRNA methylation during meiosis and interacts with all members of the yeast methyltransferase complex, indicating that it is a functional part of the MIS complex. This finding indicates that the yeast mRNA methyltransferase complex is more like that of other eukaryotes than previously thought. These findings are supported by concurrent work from Ensinck et al. (2023) that also showed that Kar4p is part of the methyltransferase complex and is required for mRNA methylation (manuscript in preparation). Previous work as well as our data show mRNA methylation via the MIS complex functions to permit entry into meiosis by regulating the transcript levels of *RME1*, encoding a key transcriptional repressor of *IME1* (Bushkin, Pincus et al. 2019). In *kar4*Δ/Δ, as in cells lacking Ime4p or expressing a catalytically dead mutant of Ime4p, we see elevated levels of Rme1p. Overexpression of *IME1* suppresses the *kar4*Δ/Δ Mei^-^ defect but does not rescue the Spo^-^ defect. *IME1* overexpression does however permit sporulation in both *ime4-cat/*Δ and *slz1*Δ/Δ. The ability of *IME1* overexpression to rescue the catalytically dead version of Ime4p and the Mei^-^ defect indicates that Kar4p’s Mei function involves mRNA methylation. The additional finding that *IME1* overexpression can suppress the defect associated with the loss of nuclear localization of the complex mediated by Slz1p indicates that Kar4p’s Mei function occurs in the nucleus. Together, these findings support a model in which Kar4p acts upstream of Ime1p and is necessary for efficient activation of *IME1* expression through nuclear mRNA methylation. In accordance with this model, we see reduced levels of Ime1p in *kar4*Δ/Δ during meiosis, and reduced levels of Rme1p suppresses the early meiotic DNA replication defect.

When the Kar4p Mei^-^ function is bypassed by *IME1* overexpression, cells progress through DNA replication and recombination, but become arrested prior to the meiotic divisions and spore formation suggesting that Kar4p has additional functions in meiosis. Work on Ime4p also suggested that the protein has additional non-catalytic functions in meiosis based on the finding that cells lacking both Rme1p and Ime4p progress through pre-meiotic S-phase, but not meiotic divisions. However, cells lacking Rme1p and expressing a catalytically dead mutant of Ime4p did progress through meiosis and form spores (Bushkin, Pincus et al. 2019). In mammalian systems, METTL3 (Ime4p) has also been shown to have non-catalytic functions that act to enhance the translation of bound transcripts (Lin, Choe et al. 2016, Wei, Huo et al. 2022). Most likely, the complex in yeast is also engaging in these additional functions.

*RIM4* overexpression can rescue the *kar4*Δ/Δ Spo^-^ defect. Given that *slz1*Δ/Δ can be fully suppressed by Ime1p overexpression alone, it is surprising that *RIM4* overexpression alone is sufficient to enhance sporulation in wild type cells and suppress *slz1*Δ/Δ. One interpretation is that suppression entails Rim4p’s early activating function rather than its later function in blocking translation. We speculate that Kar4p might work with Rim4p or other proteins to activate or support the translation of Ime2p or other regulators of genes required for the meiotic divisions and spore formation. For example, the “middle” meiotic transcription factor, Ndt80p, which regulates genes required for spore formation, is repressed by Sum1p and derepressed by Ime2p. Accordingly, the defect in spore formation may be due to mis-regulation of *NDT80* or the mis-regulation of mRNAs within *NDT80*’s regulon. This model is supported by findings presented in a companion manuscript that show reduced Ime2p levels and *NDT80* expression in *kar4*Δ/Δ with *IME1* overexpressed. The finding that the meiotic defects of both *ime4*Δ/Δ and *mum2*Δ/Δ are also suppressed by co-overexpressing both *IME1* and *RIM4* suggests that Ime4p and Mum2p may also be involved in this other function.

In summary, the genetic analysis described herein implicates *KAR4* as a major regulator of S. *cerevisiae* meiosis. It is required both early, before DNA replication and later, to facilitate entry into the meiotic divisions. These two meiotic functions are partially separable from each other, and from Kar4p’s previously characterized role in mating. This work also establishes that the yeast mRNA methylation machinery is more like that of other eukaryotes than previously thought. In a concurrent paper, we explore the impact of losing this member of the mRNA methyltransferase machinery on the meiotic transcriptome and present further evidence that the later function appears to involve mRNA methylation-independent post-transcriptional regulation during meiosis.

## Materials and Methods

### Media

In general, strains were grown in 2% glucose YPD media unless otherwise noted.

Synthetic complete (SC) media lacking the necessary nutrients for selection were made with yeast nitrogen base supplemented with the necessary nutrients except for the nutrient synthesized by the selectable marker. Pre-sporulation media was yeast nitrogen base (1% w/v), peptone (2% w/v), and potassium acetate (1% w/v). Sporulation media was either 1% (w/v) potassium acetate supplemented with histidine, uracil, and leucine or complete sporulation media which consists of yeast extract (0.25% w/v), potassium acetate (1.5% w/v), and 0.1% glucose.

### Hydroxylamine mutagenesis and genetic screen for separation-of-function alleles of *KAR4*

A centromere-based plasmid carrying *URA3* and *KAR4* with an internal 3xHA tag under the control of its native promoter (pMR2654) was mutagenized with hydroxylamine (see Table S2). The plasmid (10 µg) was incubated with 500 µl of 7% hydroxylamine solution for 20 hours at 37° (Rose, Winston et al. 1990). The reaction was stopped by adding 10 µl 5M NaCl and 50 µl 1mg/ml BSA, and the DNA was ethanol-precipitated and resuspended in TE. The mutagenized DNA was transformed into the yeast strain MY 10128 (see Table S1). Individual transformants were patched onto SC-URA media, 50 to a plate, with a positive and negative control (MY 11596 and 11597). The patches were replica-printed onto YEPD with a lawn of *MAT*α MY11297 and allowed to mate at 30° C for 3 hours (limited mating) before replica- printing on SC-LEU-LYS media to detect mating defects. The mating plates were returned to the 30° C incubator overnight, then replica printed onto SC-LEU-LYS-URA media the next day to select for diploids from all patches (overnight mating). Diploids from the overnight mating were replica printed onto Complete Sporulation media and incubated at room temperature for one day, then at 30° for three days. The sporulation plates were finally replica printed on SC-HIS-ARG media with added canavanine to detect successful entry into meiosis. The limited mating plates were compared to the meiosis plates to detect potential separation of function mutants. Potential hits were repatched (nine to a plate) from the original transformation plate and rescreened. When transformants displayed a consistent phenotype, the plasmid was extracted using a yeast mini-prep kit (Zymo Research, Irvine, CA) and electroporated into bacteria.

Bacterial colonies were grown up and the plasmids were extracted using a kit (Qiagen, Germantown, MD). The samples were sent for sequencing with *KAR4*-specific primers. After sequencing, the plasmids were retransformed into MY 10128 and assayed a final time for both mating and meiotic defects. To further address the severity of the meiotic defect, cells were sporulated as described above and spotted in 10-fold serial dilutions on the SC-HIS-ARG media. Alleles that supported wild type levels (formed colonies at the highest dilution in the series) of recombination were scored as “++++”. Alleles that only formed colonies on the 10^-2^ serial dilution were scored as “+++” and so on.

### Kar4p Structure

AlphaFold was used to generate the predicted structure of Kar4p (Jumper, Evans et al. 2021). Molecular graphics and analyses were performed with the UCSF Chimera package (http://www.rbvi.ucsf.edu/chimera<u>)/</u>, developed by the Resource for Biocomputing, Visualization, and Informatics at the University of California, San Francisco (supported by NIGMS P41-GM103311) (Pettersen, Goddard et al. 2004).

### Site-directed Mutagenesis of *KAR4*-GBD Plasmid

To move the alleles from the screen onto a *KAR4*-GBD fusion for use in yeast one-hybrid assays, we employed PCR mutagenesis. A pair of mutagenic primers was designed for each allele, and they were used to amplify the entire pMR4997 plasmid. Immediately after the PCR was finished, DpnI restriction enzyme was added to the PCR reaction tube, and the mixture was incubated at 37° for an additional 2.5 hrs. to digest the methylated template DNA. The remaining mutagenized vectors were electroporated into *E. coli*, and several candidate colonies from each plate were grown up, miniprepped, and sequenced to find the desired clones.

### Yeast One-Hybrid and Two-Hybrid

To assess whether the Kar4p alleles retained the ability to interact with Ste12p, a yeast one-hybrid assay was performed essentially as previously described (Lahav, Gammie et al. 2007). Briefly, the plasmids were transformed into the yeast two-hybrid reporter strain PJ69-4A, and dilutions of logarithmically growing cultures were normalized and spotted in 10-fold serial dilutions onto SC-TRP and SC-HIS plates with either 3 µM α-factor or an equivalent volume of methanol. The plates were incubated at 30° for several days, and interaction was assessed as ability to drive transcription of the selective marker (*HIS3*).

A two-hybrid assay was used to both verify the interaction between Kar4p and Ime4p as well as determine whether any of the Kar4p alleles impacted this interaction. *IME4* was PCR-amplified from chromosomal DNA using the primers “IME4 PGAD fwd” (5’ AAAGAGATCGAATTCCCGGGGATCCATATGATTAACGATAAACTAGT 3’) and “IME4 PGAD rev” (5’ TACTACGATTCATAGATCTCTGCAGGTTTACTGAGCAAAATATAGGT 3’). The PGAD C2 vector (pMR3659) was linearized with *BamHI*/*PstI* double restriction digest and transformed into yeast with the PCR product. Candidates growing on SC-LEU media had their plasmids extracted, sequence-confirmed (Genewiz, South Plainfield, NJ), and transformed into the PJ69-4A yeast two-hybrid reporter strain. This strain was mated to the yeast-two hybrid strain of the opposite mating type carrying the *KAR4-GBD* or *KAR4^SOFA^-GBD* constructs.

Matings were selected on SC-LEU, -TRP. Once diploids were isolated, they were induced to sporulate as described above for 24 hours before being plated in 10-fold serial dilutions on both SC-LEU, -TRP (growth), SC-HIS (selection), and SC-ADE (selection). A diploid containing the *IME4-GAD* and only an empty *GBD* was used as a negative control.

For both the one- and two-hybrid assays, alleles that facilitated growth on the selective media at every dilution were scored as “++++”. Every loss of a plus indicates that an allele only drove expression to support growth at 10-fold lower dilution. For example, an allele that could drive expression such that colonies formed in the 10^-3^ serial dilution would be scored as “++++” and an allele that could drive expression such that colonies formed in the 10^-2^ serial dilution would be scored as “+++”.

### Sporulation

Liquid cultures were sporulated as previously described (Elrod, Chen et al. 2009).

Cultures were grown overnight, back diluted into YPA, grown for 16-18 hours, washed with water, resuspended at an OD of 0.5 in 1% potassium acetate supplemented with the amino acids necessary for each strain, and incubated at 26°C for varying amounts of time depending on the experiment being conducted. For sporulation assays involving the overexpression of *IME1* and *RIM4*, the only difference was that these cultures were induced with 1 µM of β-estradiol or an equivalent volume of 100% ethanol upon addition of the cells to the 1% potassium acetate media. Experiments using strains from the SK1 background were sporulated in much the same way with the only differences being cells were resuspended in 1% potassium acetate at an OD of 2.0 and were sporulated at 30°C.

### mRNA Methylation

SK1 cultures were sporulated for four hours as described above. Cells were lysed using the FastPrep (MP) with four rounds of bead beating at 4.0 m/s for 40 seconds with one minute on ice in between each round. Total RNA was then harvested using the Qiagen RNeasy kit. mRNA was purified from these total RNA samples using Oligod(T) magnetic beads (Thermofisher). mRNA methylation was measured using the EpiQuik fluorometric m^6^A RNA methylation quantification kit (EpiGenTek). Briefly, purified mRNA was bound to wells and then washed before the addition of an antibody to m^6^A. Wells were then washed again before the addition of a detection antibody followed by an enhancer/developer solution. The fluorescence was measured using a plate reader at 530EX/590EM nm and m^6^A levels were quantified for each sample using a standard curve generated from a positive control provided with the kit. Measurements from an *ime4*Δ/Δ strain were used to subtract background from the data collected from other mutants.

### Protein Extraction

Proteins were extracted using alkaline lysis followed by TCA precipitation. Briefly, samples were incubated with 150 µl of 1.85 M NaOH on ice for 10 minutes. An equal volume of 50% TCA was then added to the samples and they were left to incubate at 4°C for 10 minutes.

Samples were then washed with acetone before being resuspended in 100 µl of 2x sample buffer. Samples were then boiled for five minutes and then kept on ice for five minutes before being run on an SDS-PAGE gel or stored for later use at -80°C.

### Western Blotting

Protein samples were resolved on an appropriate percentage SDS-PAGE gel before being transferred to a PVDF membrane using a semi-dry transfer apparatus (TransBlot SD BioRad) at 16 volts for 36 minutes. After transfer, membranes were blocked for 30 minutes using 10% milk in TBS before being incubated with primary antibody (anti-HA (12CA5) 1:1,000, anti-HA (rabbit: Cell Signaling C29F4) 1:2,500, anti-MYC (rabbit: Novus NB600-336SS) 1:1,000, anti-FLAG (Sigma M2) 1:1000, anti-GFP (BD Living Colors 632377) 1:1000) for one hour at room temperature. Membranes were then washed three times for ten minutes each with 0.1% TBS- Tween20 (TBST) before being incubated with a secondary antibody in 1% milk in TBST for 30 minutes at room temperature. Incubation with secondary antibody (Donkey anti-mouse IgG (Jackson ImmunoResearch) 1:10,000 or Donkey anti-rabbit IgG (ImmunoResearch) 1:20,000) was then followed by another three washes for ten minutes each with TBST before being incubated with Immobilon Western HRP substrate (Millipore) for 5 minutes and imaged using the G-Box from SynGene. Densitometry was conducted using ImageJ.

### Co-Immunoprecipitation

Cells were sporulated for four hours as described above and a volume of cells equivalent to 50 OD units was harvested for each strain. Cells were resuspended in a lysis buffer (1% Triton-X 100, 0.2 M Tris-HCl pH 7.4, 0.3 M NaCl, 20% glycerol, 0.002 M EDTA) supplemented with 10x protease inhibitor (Pierce) and lysed using the FastPrep (MP) as described above. Protein extracts were then incubated with 15 µL of anti-MYC or HA magnetic beads (Pierce) for one hour at room temperature. Beads were washed three times with lysis buffer before being resuspended in 2x sample buffer and boiled for 5 minutes to elute proteins off the beads. Samples were then resolved on 8% SDS-PAGE gels and western blotting was conducted as described above.

### Microscopy

For microscopy, 100 µl of a sporulated culture expressing GFP-*TUB1* integrated at *TUB1* (Straight, Marshall et al. 1997) and *SPC42*-mCherry was spun down and resuspended in 10 µl of 1% potassium acetate media. Cells were imaged on a DeltaVision deconvolution microscope (Applied Precision, Issaquah, WA) based on a Nikon TE200 (Melville, NY) using a 100x/numerical aperture 1.4 objective, a 50W mercury lamp, and a Photometrics Cool Snap HQ charge-coupled device camera (Photometrics, Tucson, AZ). All images were deconvolved using Applied Precision SoftWoRx imaging software. At least one hundred cells were counted in all cases, and the percentage of cells in each meiotic stage including mature spores were determined using the morphology of the spindle and number of spindle pole bodies.

### Flow Cytometry

Cells were induced to sporulate as described above. At each time point measured, ∼5x10^6^ cells were pelleted and washed in dH2O. Cells were fixed in 70% ethanol and stored at -20 °C for one hour up to several days. After fixation, cells were washed in 500 µl 50mM sodium citrate (pH 7.2) and incubated in 0.5 ml sodium citrate containing 0.25 mg/ml RNaseA for two hours at 37°C. After RNaseA treatment, 500 µl of 8 µg/mL propidium iodide in sodium citrate buffer was added to each sample and incubated overnight in the dark at 4 °C. The cells were then sonicated at 50% duty cycle, output setting 3, for 4x12 pulses on ice before being transferred to Falcon 2054 tubes. FACS was conducted at the Flow Cytometry and Cell Sorting Shared Resource core at Georgetown University and data was analyzed using FCS Express.

### High-copy suppression assays and screen

To find suppressors of *KAR4*’s “Mei” function, MY10128 was transformed with a YEp24 2µ plasmid library (Carlson and Botstein 1982) and the colonies were mated overnight against a lawn of MY11297. Diploids from this cross were selected with SC-LEU-LYS-URA media and then replica-printed onto complete sporulation media (described above) and incubated at room temperature for one day, then at 30°C for three days. The sporulation plates were finally replica- printed on SC-His-Arg media with added canavanine to detect successful recombination/suppression. When colonies grew after meiosis, plasmids were extracted from them and sent for sequencing. *IME1* and *IME2* were subcloned into pRS426 to confirm that they were the causal genes on the isolated fragments.

For the screen to find genes that rescued the sporulation defect of Z4EV-*IME1* induced *kar4*Δ diploids, the same 2µ library was transformed into the Z4EV-*IME1 kar4*Δ diploid strain. Transformants were washed off plates and induced to sporulate by addition of 1 µM estradiol. After 3 days at 30° C, the cultures were digested with zymolyase to kill diploid cells, and dilutions were plated on SC –URA media to retain the plasmids in any surviving cells or spores. Candidate colonies were replica-plated onto a *sst2*Δ lawn. Cells that formed halos on the *sst2*Δ lawn were further tested for papillation on media containing canavanine to eliminate any diploids that survived zymolyase digestion. Plasmids were extracted from haploid cells and sent for sequencing, and any plasmids containing fragments other than *KAR4* were retransformed and reinduced to confirm that they led to spore formation. *RIM4* was then subcloned into pRS426.

### PZ4EV-*IME1* and PZ3EV-*RIM4* Construction

The β-estradiol-inducible promoter (PZ4EV) tagged with KanMX was PCR-amplified from plasmid pRSM29 (gift of R. Scott McIsaac) using *IME1*-specific promoters. The PCR product was transformed into MS3215, and transformants were selected based on G418 resistance and confirmed by colony PCR. A successful integrant was mated against strain DBY12416 (gift of the Botstein laboratory), which contains the gene for the *Z4EV* artificial transcription factor under the *ACT1* promoter. Spores from this cross were dissected to yield a haploid strain that contained *kar4*Δ, PZ4EV-*IME1*, *ura3*Δ and *Z4EV*. The PZ3EV-*RIM4* strain was made basically the same way except the promoter was amplified from vector RB3483 (gift of Patrick Gibney) and made in a haploid strain containing the appropriate *Z3EV* artificial transcription factor.

### qPCR

Approximately 10 OD units of cells were harvested at the indicated time points. RNA was extracted from cells using the Qiagen RNeasy kit. Briefly, cells were lysed, and the supernatant was cleared and mixed with an equal volume of 70% ethanol. The mixture was then loaded on the column provided with the kit and the wash steps were followed as detailed in the kit’s protocol. The on-column DNase treatment (Qiagen) was used to remove any DNA from the RNA sample. Total RNA was eluted in 100 µl of RNase free water. cDNA libraries were constructed using the High-Capacity cDNA Reverse Transcription kit (Applied Biosystems) with 10 µl of the total RNA sample. The concentration of the resulting cDNA was measured using a nanodrop. qPCR reactions were set up using Power SYBR Green PCR Master Mix (Applied Biosystems) with a volume of the total RNA sample equivalent to 50 ng. The reactions were run on a CFX96 Real-Time System (BioRad) with reaction settings exactly as described in the master mix instructions with the only change being the addition of a melt curve at the end of the program. Results were analyzed using CFX Maestro. Primer sequences are as follows: *PGK1* Forward 5’-CTCACTCTTCTATGGTCGCTTTC-3’, *PGK1* Reverse 5’- AATGGTCTGGTTGGGTTCTC-3’, *IME1* Forward 5’-ATGGCAACTGGTCCTGAAAG-3’, and *IME1* Reverse 5’-GGAACGTAGATGCGGATTCAT-3’.

## Acknowledgements

We thank Anne Rosenwald and Luke Berchowitz for helpful feedback on this manuscript and project as well as members of the Rose Lab. We thank May Husseini for making media and helping with initial mutant screen. We thank Folkert van Werven for providing the plasmid for tagging *IME1* and for sharing information prior to publication. This work was supported by NIH grants GM037739 and GM126998 to MDR.

---

**S Fig 1.**
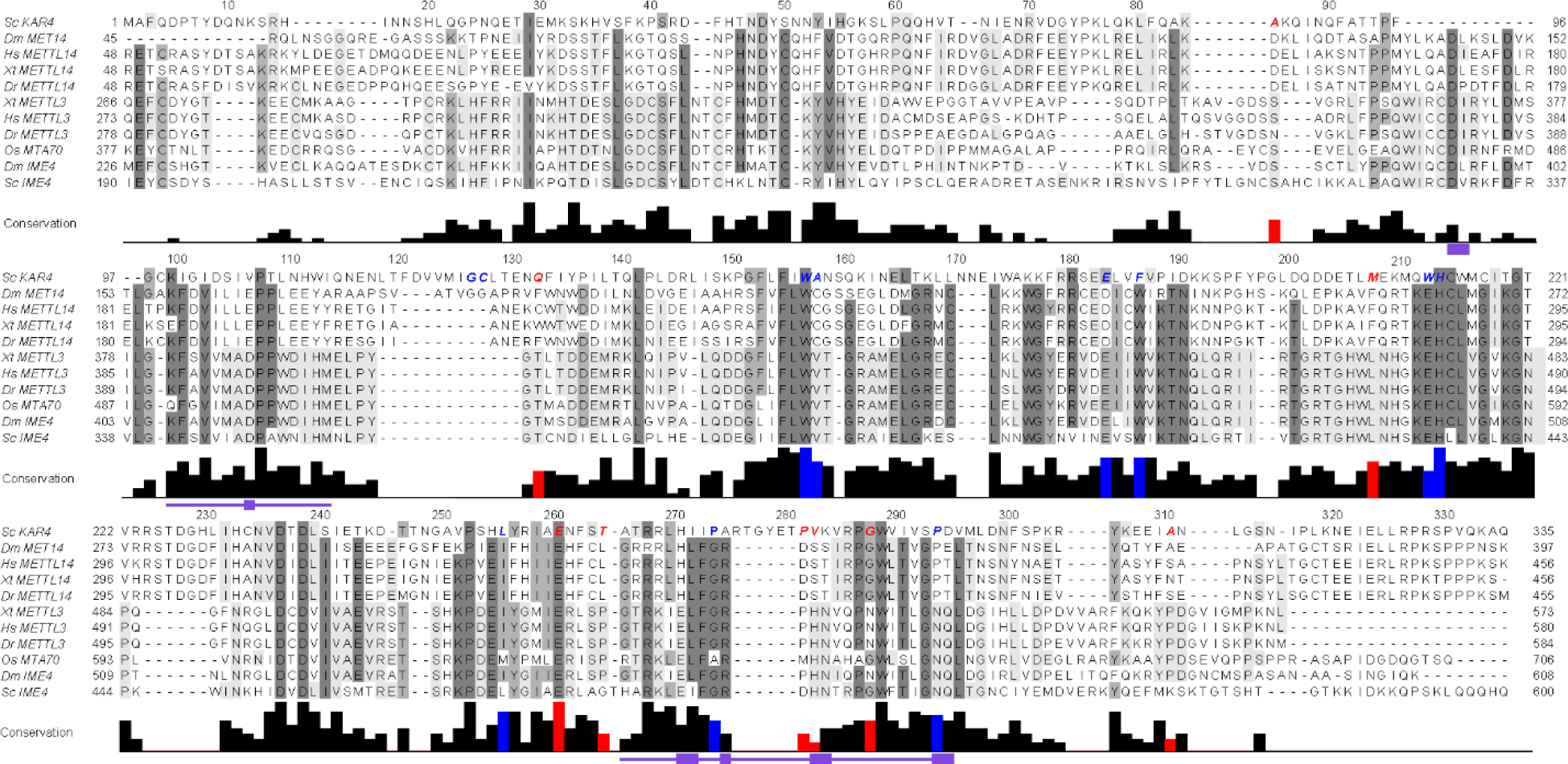
Conservation of *KAR4* and the separation of function alleles across the KAR4 and ***IME4* protein lineages.** Sequence conservation of *KAR4* and *IME4* and a broad set of orthologs from a range of vertebrates, invertebrates, and plants: Sc, *Saccharomyces cerevisiae*; Dm, *Drosophila melanogaster*; Hs, Homo sapiens; Dr, *Danio rerio*; Xt, *Xenopus tropicalis*; Os; *Oryza sativa*. The degree of conservation is indicated by the height of the histogram below the sequence and by the shade of the background. Note the presence of blocks that are specific to Kar4p or Ime4p orthologues (light grey), as well as regions conserved between both paralogs (dark grey). Alleles defective for mating are in blue and alleles defective for meiosis are in red. Functional motifs associated with the methyltransferase activity of Ime4p orthologues (Bujnicki et al. 2002) are indicated by the purple regions below the sequence.

**S Fig 2.**
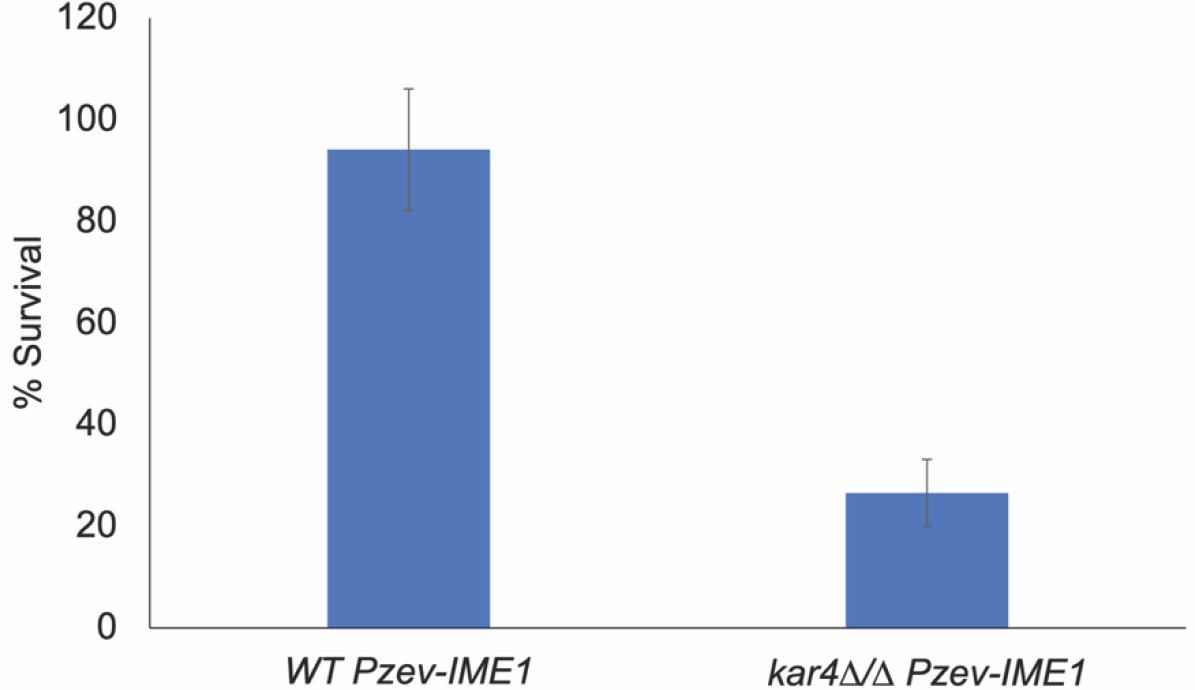
*kar4*Δ/Δ mutants lose viability after *IME1* overexpression. Viability of *kar4*Δ/Δ mutants 48 hours after induction of *IME1* expression in sporulation conditions. Viability was assessed as the change in colony forming units between t = 0 (before *IME1* induction) and t = 48 hours after *IME1* induction. Error bars represent the standard deviation of three biological replicates.

**S Fig 3.**
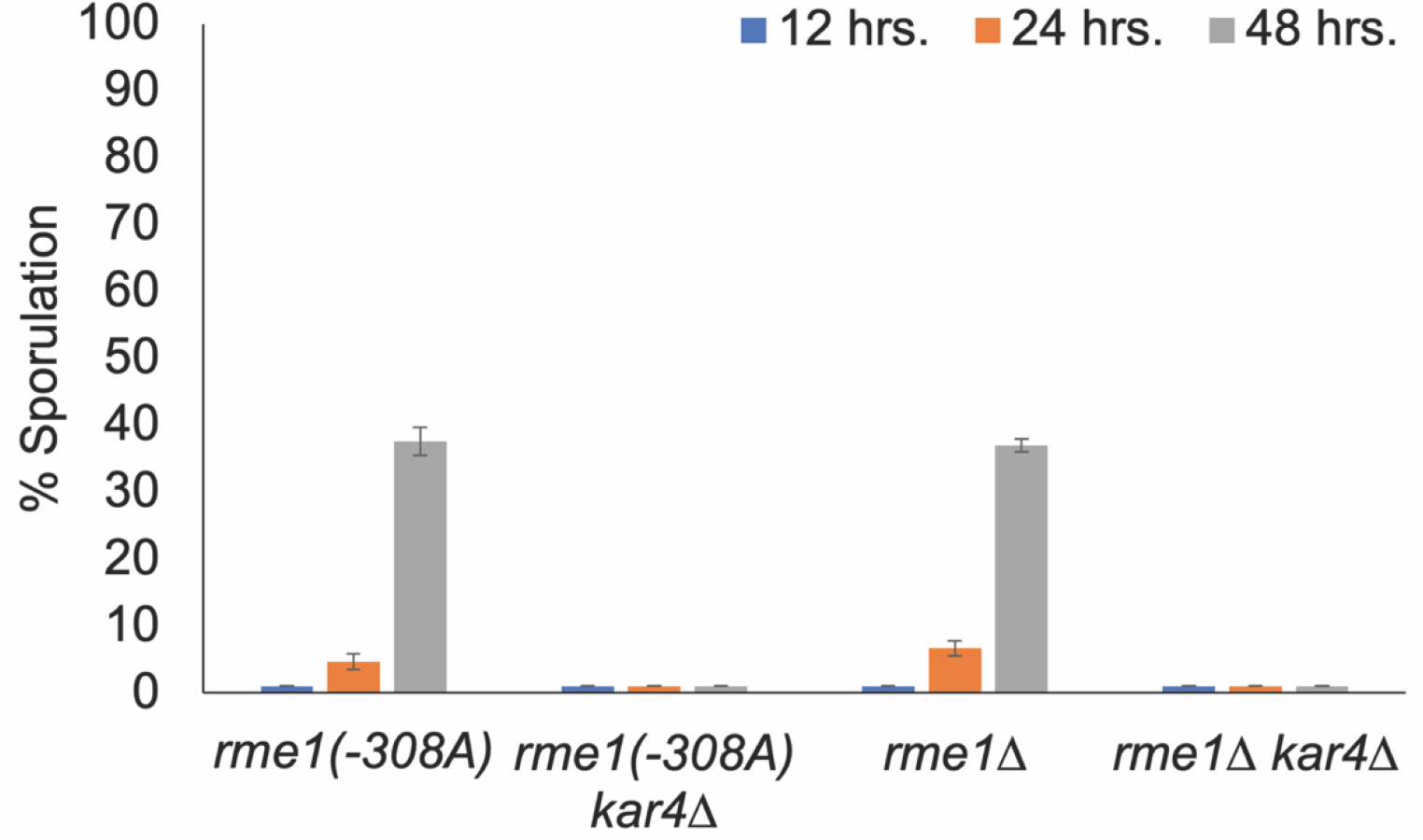
*kar4*Δ*rme1*Δ/ *kar4*Δ*rme1*Δ and *kar4*Δ*rme1(-308A)*/ *kar4*Δ*rme1(*-308A) mutants do not sporulate. Spores were counted at the indicated time points in *kar4*Δ*rme1*Δ/ *kar4*Δ*rme1*Δ, *kar4*Δ*rme1(- 308A)*/ *kar4*Δ*rme1(*-308A), and the single mutants of each *RME1* allele. All dyads, triads, and tetrads were counted. At least 100 cells were counted for each time point. Error bars represent the standard deviation of three biological replicates.

**S Fig 4.**
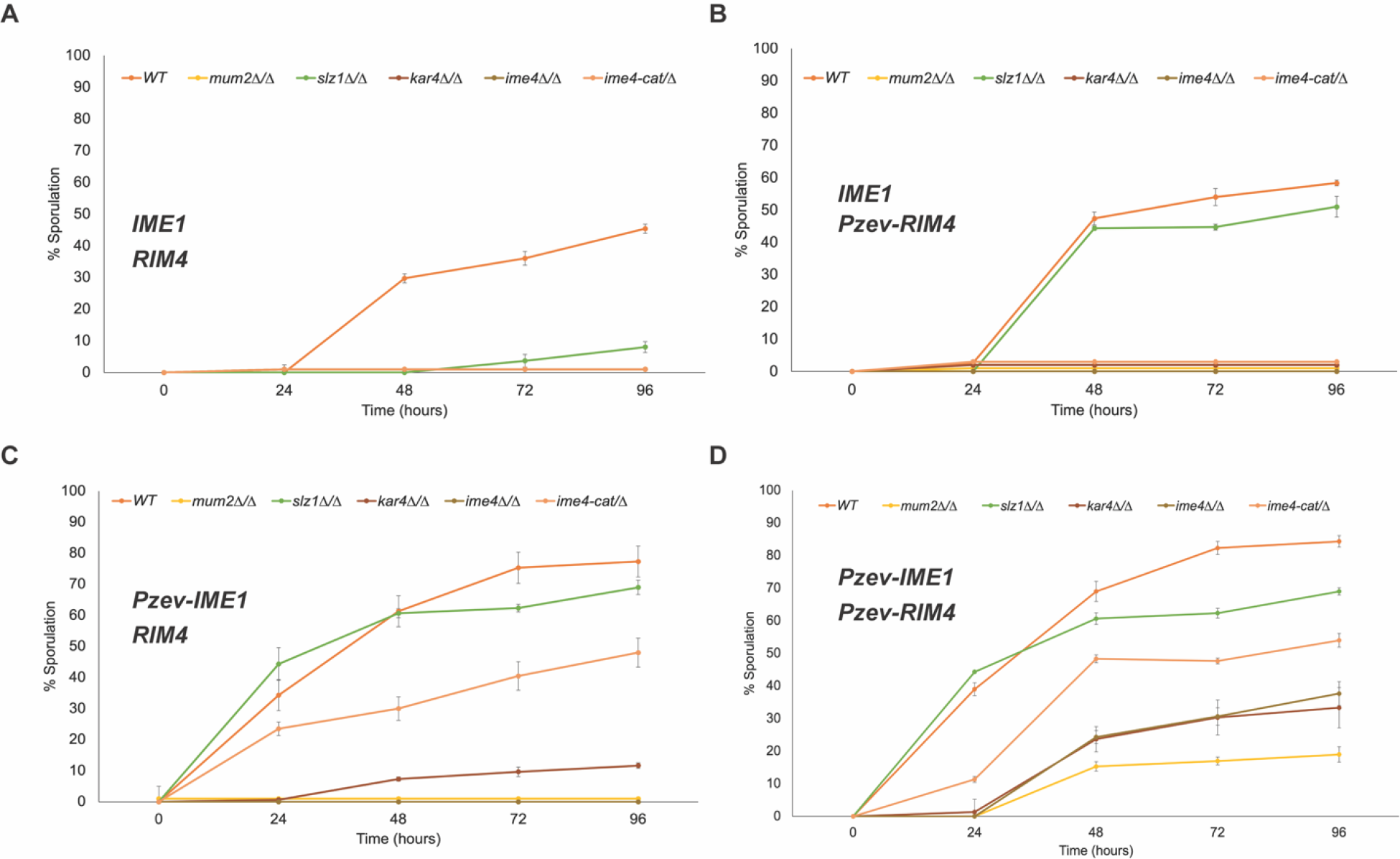
Sporulation of methyltransferase complex mutants across an extended time course **of meiosis.** (A) Spore counts for *IME1/RIM4* in the indicated mutants. (B) Spore counts for *IME1/PzevRIM4* in the indicated mutants. (C) Spore counts for *PzevIME1/RIM4* in the indicated mutants. (D) Spore counts for *PzevIME1/PzevRIM4* in the indicated mutants. For each experiment all dyads, triads, and tetrads were counted at the indicated times. For experiments involving overexpression (B, C, and D), 1 µM of estradiol was used to induce expression. At least 100 cells were counted for each time point.

**Table S1.** Strains used for this study. All auxotrophic markers are standard BY alleles.

| Strain Name | Genotype | Strain | Reference |
| --- | --- | --- | --- |
| MY 10128 | *leu2 his3 ura3 met15 kar4::KanMX* | S288c |  |
| MY 11297 | *ura3 leu2 his3 lys2 can1::LEU2+-MFA1pr-HIS3 kar4::KanMX* | S288c |  |
| MY 16294 | *URA3::GFP-TUB1/+ SPC42-mcherry::HIS3/+ +/kar4::URA3 HPH::Pz3ev-IME1/+ leu2::ACT1PR-3ZEV-NatMX/leu2 his3/+ ura3/" met15/+* | S288c |  |
| MY 16295 | *URA3::GFP-TUB1/+ SPC42-mcherry::HIS3/+ kar4::KANMX/kar4::URA3 HPH::Pz3ev-IME1/+ leu2::ACT1PR-3ZEV-NatMX/leu2 his3/+ ura3/" met15/+* | S288c |  |
| PJ69-4A | *trp1-901 leu2-3,112 ura3-52 his3-d200 gal4 gal80 GAL2-ADE2 LYS2::GAL1::HIS3 met2::GAL7-lacZ* | S288c | James et al. 1996 |
| PJ69-4@ | *trp1-901 leu2-3,112 ura3-52 his3-d200 gal4 gal80 GAL-ADE2 LYS2::GAL1::HIS3 met2::GAL7-lacZ* | S288c | James et al. 1996 |
| MY 16325 | *ura3/" lys2/" ho::LYS2/" arg4-BgIII/+ leu2::HisG/+* | SK1 |  |
| MY 16326 | *ura3/" lys2/" ho::LYS2/" arg4-BgIII/+ leu2::HisG/+ ime4::URA3/"* | SK1 |  |
| MY 16351 | *ho::LYS2/" arg4-BgIII/+ +/leu2::hisG lys2-SK1/" ura3::hisG/" kar4::KANMX/+* | SK1 |  |
| MY 16353 | *ho::LYS2/" arg4-BgIII/+ +/leu2::hisG lys2-SK1/" ura3::hisG/" slz1::NATMX/"* | SK1 |  |
| MY 16356 | *ho::LYS2/" arg4-BgIII/+ +/leu2::hisG lys2-SK1/" ura3::hisG/" kar4::KANMX/" slz1::NATMX/"* | SK1 |  |
| MY 16256 | *leu2/" his3/" ura3/" lys2/+ met15/+ kar4::KANMX/" MUM2-4MYC::HIS3/+ [URA+]* | S288c |  |
| MY 16257 | *leu2/" his3/" ura3/" lys2/+ met15/+ kar4::KANMX/" MUM2-4MYC::HIS3 [KAR4-HA-URA]* | S288c |  |
| MY 16405 | *SLZ1-3HA::HIS3/+ KAR4-9MYC::KANMX/+ kfc1::KANMX/+ his3/" met15/+ leu2/" ura3/" lys2/+* | S288c |  |
| MY 16409 | *SLZ1-3HA::HIS3/+ his3/" met15/+ leu2/" ura3/" lys2/+* | S288c |  |
| MY 13813 | *KANMX::Pzev4-IME1/ime1::KANMX kar4::KANMX/" ura3d52/" his3d100/+ leu2::ACT1PR-42EV-NATMX/leu2 ybr032w::LEU2/+* | S288c |  |
| MY 13815 | *KANMX::Pzev4-IME1/ime1::KANMX kar4::KANMX/+ ura3d52/" his3d100/+ leu2::ACT1PR-42EV-NATMX/leu2 ybr032w::LEU2/+* | S288c |  |
| MY 16456 | *rme1::KANMX/" leu2/" his3/" ura3/" lys2/+ met15/+* | S288c |  |
| MY 16557 | *his3/" leu2/" ura3/" lys2/+ met15/+ rme1(-308A)::KANMX/"* | S288c |  |
| MY 16559 | *his3/" leu2/" ura3/" lys2/+ met15/+ rme1(-308A)::KANMX/" kar4::HPHMX/"* | S288c |  |
| MY 16566 | *leu2/" his3/" ura3/" met15/+ lys2/+ rme1::KANMX/" kar4::HPHMX/"* | S288c |  |
| MY 16550 | *his3/" leu2/" ura3/" lys2/+ met15/+ GFP-IME1/"* | S288c |  |
| MY 16570 | *his3/" leu2/" ura3/" lys2/+ met15/+ GFP-IME1/" kar4::HPHMX/"* | S288c |  |
| MY 16622 | *his3/" leu2/" ura3/" lys2/+ met15/+ rme1Δ::KANMX/" kar4Δ::HPHMX/" GFP-IME1/"* | S288c |  |
| MY 16563 | *leu2/" his3/" ura3/" met15/+ lys2/+ 3xFLAG-RME1/"* | S288c |  |
| MY 16569 | *leu2/" his3/" ura3/" met15/+ lys2/+ 3xFLAG-RME1/" kar4::HPHMX/"* | S288c |  |
| SAy914 | *lys2/" ho::LYS2/" 3xmyc-IME4/"* | SK1 | Agarwala et al. 2012 |
| MY 16543 | *lys2/" ho::LYS2/" 3xmyc-IME4/" kar4::HPHMX/kar4::KANMX* | SK1 |  |
| MY<br>16616 | *his3/" leu2/leu2::pACT1-Z3EV-natMX +/lys2 met15/+ ura3/"* | S288c |  |
| MY<br>16532 | *HPHMX::PZ3EV-RIM4/+ leu2::act1pr-Z3EV-NATMX/leu2 ura3/" his3/+ lys2/+* | S288c |  |
| MY<br>16533 | *HPHMX::PZ3EV-IME1/+ HPH::PZ3EV-RIM4/+ leu2Δ0::act1pr-Z3EV-NATMX/" ura3/"* | S288c |  |
| MY<br>16534 | *HPHMX::PZ3EV-IME1/+ leu2::act1pr-Z3EV-NATMX/leu2 ura3/" his3/+ lys2/+* | S288c |  |
| MY<br>16617 | *his3/" leu2/leu2::pACT1-Z3EV-NATMX +/lys2 met15/+ ura3/" kar4::KANMX/"* | S288c |  |
| MY<br>16531 | *HPHMX::PZ3EV-IME1/+ leu2::act1pr-Z3EV-NATMX/leu2 ura3/" his3/+ met15/+ kar4::KANMX/kar4::URA3* | S288c |  |
| MY<br>16535 | *HPHMX::PZ3EV-RIM4/+ leu2::act1pr-Z3EV-NATMX/leu2 ura3/" kar4::LEU2/kar4::KANMX4 lys2/+* | S288c |  |
| MY<br>16536 | *HPHMX::PZ3EV-IME1/+ HPHMX::PZ3EV-RIM4/+ leu2::act1pr-Z3EV-NATMX/" ura3/" kar4::LEU2/kar4::URA3* | S288c |  |
| MY<br>16619 | *his3/" leu2/leu2::pACT1-Z3EV-NATMX +/lys2 met15/+ ura3/" mum2::KANMX/"* | S288c |  |
| MY<br>16525 | *HPHMX::PZ3EV-IME1/+ leu2::act1pr-Z3EV-NATMX/leu2 ura3/" lys2/+ his3/+ mum2::URA3/mum2::KANMX* | S288c |  |
| MY<br>16526 | *HPHMX::PZ3EV-RIM4/+ leu2::act1pr-Z3EV-NATMX/leu2 ura3/" lys2/+ his3/+ mum2::LEU2/mum2::KANMX* | S288c |  |
| MY<br>16527 | *HPHMX::PZ3EV-IME1/+ HPHMX::PZ3EV-RIM4/+ leu2::act1pr-Z3EV-NATMX/" ura3/" mum2::URA3/mum2::LEU2* | S288c |  |
| MY<br>16620 | *his3/" leu2/leu2::pACT1-Z3EV-NATMX +/lys2 met15/+ ura3/" slz1::KANMX/"* | S288c |  |
| MY<br>16537 | *HPHMX::PZ3EV-RIM4/+ leu2::act1pr-Z3EV-NATMX/leu2 ura3/" slz1::LEU2/slz1::KANMX his3/+ met15/+* | S288c |  |
| MY<br>16538 | *HPHMX::PZ3EV-IME1/+ leu2::act1pr-Z3EV-NATMX/leu2 ura3/" slz1::URA3/slz1::KANMX his3/+ lys2/+* | S288c |  |
| MY<br>16539 | *HPHMX::PZ3EV-IME1/+ HPHMX::PZ3EV-RIM4/+ leu2::act1pr-Z3EV-NATMX/" ura3/" slz1::URA3/slz1::LEU2* | S288c |  |
| MY<br>16631 | *his3/" leu2/leu2::pACT1-Z3EV-NATMX +/lys2 met15/+ ura3/" ime4::KANMX/"* | S288c |  |
| MY<br>16434 | *HPHMX::PZ3EV-RIM4/+ leu2::act1pr-Z3EV-NATMX/leu2 ura3/" met15/+ his3/+ ime4::LEU2/ime4::URA3* | S288c |  |
| MY<br>16435 | *HPHMX::PZ3EV-IME1/+ HPHMX::PZ3EV-RIM4/+ leu2::act1pr-Z3EV-NATMX/" ura3/" ime4::LEU2/ime4::URA3* | S288c |  |
| MY<br>16313 | *HPHMX::PZ3EV-IME1/+ leu2::act1pr-Z3EV-NATMX/leu2 ura3/" met15/+ his3/+ ime4::LEU2/ ime4::URA3* | S288c |  |
| MY<br>16621 | *his3/" leu2/leu2::pACT1-Z3EV-NATMX +/lys2 met15/+ ura3/" ime4-cat/"* | S288c |  |
| MY<br>16615 | *his3/" leu2::pACT1-Z3EV-NATMX/leu2 ura3/" HPHMX::PZ3EV-IME1/+ ime4::LEU2/ime4-cat HPHMX::PZ3EV-RIM4/+* | S288c |  |
| MY<br>16506 | *HPHMX::PZ3EV-IME1/+ ime4::URA3/ime4-cat his3/+ leu2/leu2::act1pr-Z3EV-NATMX lys2/+ ura3/"* | S288c |  |
| MY<br>16507 | *his3/+ lys2/+ ura3/" ime4-cat/ime4::LEU2 HPHMX::PZ3EV-RIM4/+ leu2::act1pr-Z3EV-NATMX/leu2* | S288c |  |

**Table S2.**
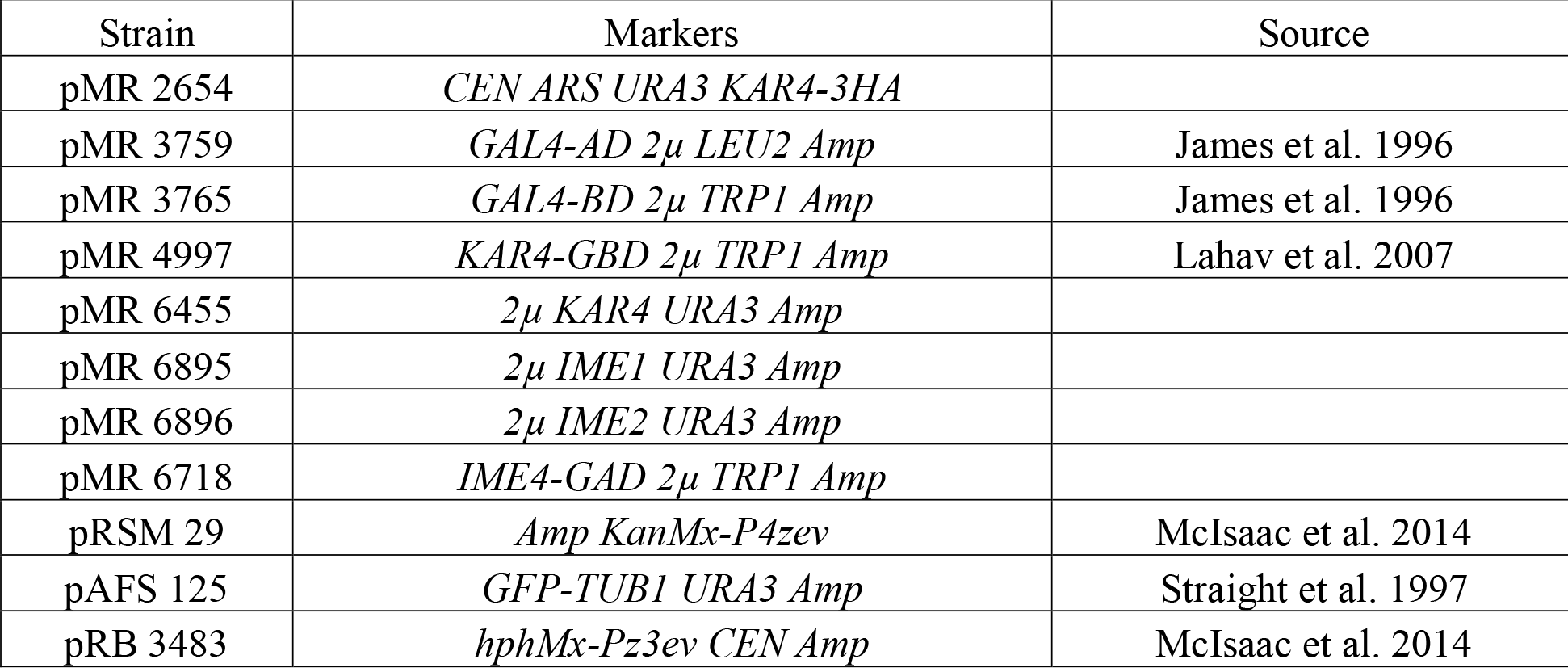
Plasmids used in this paper.

| Strain | Markers | Source |
| --- | --- | --- |
| pMR 2654 | *CEN ARS URA3 KAR4-3HA* |  |
| pMR 3759 | *GAL4-AD 2μ LEU2 Amp* | James et al. 1996 |
| pMR 3765 | *GAL4-BD 2μ TRP1 Amp* | James et al. 1996 |
| pMR 4997 | *KAR4-GBD 2μ TRP1 Amp* | Lahav et al. 2007 |
| pMR 6455 | *2μ KAR4 URA3 Amp* |  |
| pMR 6895 | *2μ IME1 URA3 Amp* |  |
| pMR 6896 | *2μ IME2 URA3 Amp* |  |
| pMR 6718 | *IME4-GAD 2μ TRP1 Amp* |  |
| pRSM 29 | *Amp KanMx-P4zev* | McIsaac et al. 2014 |
| pAFS 125 | *GFP-TUB1 URA3 Amp* | Straight et al. 1997 |
| pRB 3483 | *hphMx-Pz3ev CEN Amp* | McIsaac et al. 2014 |

